# Base editing in *Streptomyces* with Cas9-deaminase fusions

**DOI:** 10.1101/630137

**Authors:** Zhiyu Zhong, Junhong Guo, Liang Deng, Li Chen, Jian Wang, Sicong Li, Wei Xu, Zixin Deng, Yuhui Sun

**Affiliations:** Key Laboratory of Combinatorial Biosynthesis and Drug Discovery, Ministry of Education, and Wuhan University School of Pharmaceutical Sciences, Wuhan 430071, People’s Republic of China

## Abstract

Conventional CRISPR/Cas genetic manipulation has been profitably applied to the genus *Streptomyces*, the most prolific bacterial producers of antibiotics. However, its reliance on DNA double-strand break (DSB) formation leads to unacceptably low yields of desired recombinants. We have adapted for *Streptomyces* recently-introduced cytidine base editors (CBEs) and adenine base editors (ABEs) which enable targeted C-to-T or A-to-G nucleotide substitutions, respectively, bypassing DSB and the need for a repair template. We report successful genome editing in *Streptomyces* at frequencies of around 50% using defective Cas9-guided base editors and up to 100% by using nicked Cas9-guided base editors. Furthermore, we demonstrate the multiplex genome editing potential of the nicked Cas9-guided base editor BE3 by programmed mutation of nine target genes simultaneously. Use of the high-fidelity version of BE3 (HF-BE3) essentially improved editing specificity. Collectively, this work provides a powerful new tool for genome editing in *Streptomyces*.

## Introduction

Streptomycetes are high G+C Gram-positive filamentous bacteria that are extremely abundant in nature. They have been intensively studied since the 1960s both for their intriguing developmental biology and for their extraordinary ability to synthesize numerous bioactive secondary metabolites^1,2^. However, the identification and engineering of the gene clusters governing the biosynthetic pathway present an enormous challenge due to the lack of convenient and efficient genome editing tools.

The clustered regularly interspaced short palindromic repeat (CRISPR)/CRISPR-associated protein (Cas) system is a breakthrough technology that has been developed rapidly in recent years and widely applied in many species, including *Streptomyces*^3–8^. It has accelerated genetic manipulation in *Streptomyces* by introducing a precise DNA double-strand break (DSB) at the single guide RNA (sgRNA) target site and subsequently repairing it through non-homologous end joining (NHEJ) or homologous recombination (HR), resulting in either stochastic insertions/deletions or precise genome editing around the DSB. However, the actual editing capacity *in situ* depends on the efficiency of the intrinsic or reconstituted repair machinery, which in contrast to eukaryotes is inefficient in bacteria^9^ and may vary among different species, hampering the application of this system. In particular, current technology cannot easily deliver the desirable goal of multiple simultaneous precise editing.

More recently, a new technology has emerged from CRISPR/Cas termed “base editing (BE)” in which the fusion of a cytidine deaminase to a Cas9 variant enables programmable C•G → T•A base-pair conversion without the need for DSB formation or template donor DNA^10,11^ (Fig. 1a). In its initial formulation the base editor (BE1) tethers a single-stranded DNA-specific cytidine deaminase (APOBEC1) to the N-terminus of a catalytically defective Cas9 (dCas9). *In vitro* experiments have revealed that BE1 catalyzes a C-to-U deamination reaction within the 4^th^-8^th^ nucleotides of the protospacer. The resulting U is read as a T in the process of DNA repair or replication, thereby effecting a permanent C•G → T•A base-pair conversion on genomic DNA. A Uracil glycosylase inhibitor (UGI) is incorporated in the second-generation base editor (BE2) which protects the generated U bases from excision by uracil DNA glycosylase (UDG), thus enhancing the desired C-to-T editing in living cells. Further replacement of dCas9 by a Cas9 nickase (nCas9) in the third-generation base editor (BE3) enables a nick to be generated on the non-edited strand. Mismatch repair (MMR) is induced by the nick in mammalian cells, thereby converting a U•G mismatch into U•A pair more effectively. Owing to its unique mechanism, the base editing system overcomes the drawbacks of DSB-based genome editing approaches. To date, a series of expansions and improvements on the methodology have been performed by multiple studies^12–16^, and these base editors have been applied in a wide variety of animals^17–19^, plants^20,21^ and bacteria^22–27^. In addition to the cytidine base editors (CBEs), adenine base editors (ABEs) using a heterodimer of wildtype and engineered adenosine deaminase (TadA-TadA*) have been developed that extend editing to allow the A•T → G•C base-pair conversion^28^ in various organisms^19,29–32^.

**Fig. 1.**
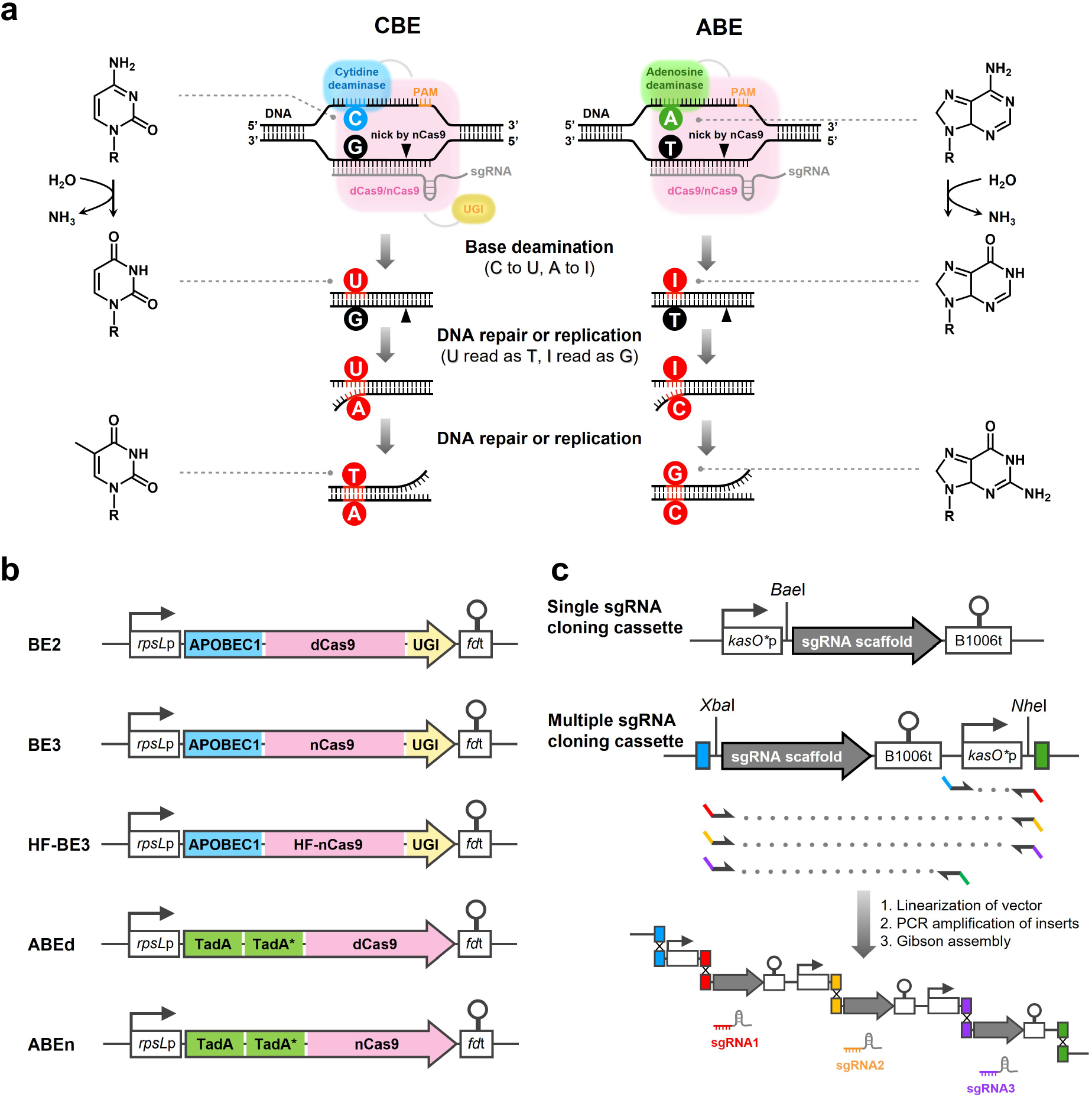
Representation of the base editing strategy and basic construct of the *Streptomyces* base editing system. **a** Procedure for CBE- and ABE-mediated base editing. dCas9, catalytically defective Cas9 with D10A and H840A mutation. nCas9, Cas9 nickase with D10A mutation. PAM, protospacer adjacent motif. UGI, uracil glycosylase inhibitor. The Cas9:sgRNA:DNA complex provides single strand DNA substrate for the deaminase. The chemistry of the base conversions is illustrated on both sides, similarities between the intermediate bases and the ultimate bases enables the conversions to take place. In contrast to dCas9, nCas9 nicks the non-edited strand to make U•G mismatch conversion into U•A or I•T mismatch conversion into I•C more effective. UGI protects the generated U bases from being excised by endogenous uracil DNA glucosylase. **b** Schematic of the base editors tested in this study. *rps*Lp, *rps*L promoter. *fd*t, fd terminator. ABPOBEC1, cytidine deaminase. TadA and TadA*, wildtype and engineered adenosine deaminase. **c** Schematic of the sgRNA cloning strategy. *kas*O*p, *kasO\** promoter. B1006t, B1006 terminator.

Base substitutions in the codons of the open reading frames (ORFs) can be used to programme amino acid substitutions (Supplementary Fig. 1 and Supplementary Table 1). It thus offers a powerful means to engineer industrial antibiotic-producing strains. For example, among the repertoire of amino acid-coding substitutions generated by CBE-mediated C•G → T•A editing, conversions of four codons into stop codons serve as a convenient alternative strategy for gene disruption. ABE-mediated A•T → G•C base-pair conversion provides additional flexibility for genome editing, and can in principle reverse mutations generated by CBE if the editing window permits.

Here, we present the use of CBEs and ABEs to create targeted base substitutions in the model strain *Streptomyces coelicolor* M145 at up to 100% efficiency. The remarkable potential of this system is demonstrated by simultaneous disruptions of nine different polyketide synthase (PKS) gene clusters in the industrially-important strain *Streptomyces avermitilis* MA-4680.

## Results

### Design of base editing systems in *Streptomyces*

Although the nCas9-guided base editors have proved more efficient than dCas9-guided base editors in eukaryotic cells^10^, it is unknown whether nickase activity to produce single strand breaks (SSB) is harmful to the *Streptomyces* cell. Therefore, CBE and ABE systems were tested with both dCas9 and nCas9 versions (Fig. 1b). Given the existence of highly-active UDGs in the *Streptomyces*, UGI was also adapted in the tested CBEs to protect the uracil-containing intermediate (Fig. 1b). The base editors were codon optimized for *Streptomyces* to promote their expression in these typically high G+C content genomes. A shuttle vector that allows for conjugal transfer from the donor *Escherichia coli* into *Streptomyces* was used to introduce the BE system (Supplementary Fig. 2). For sgRNA cloning, in addition to the traditional single sgRNA cloning cassette (SSCC) that uses a type-IIS restriction endonuclease site to insert the 20-nt spacer, we incorporated a multiple sgRNA cloning cassette (MSCC) that adapts the 20-nt spacers as homologous overlaps for Gibson assembly to enable convenient single or multiple sgRNA cloning (Fig. 1c and Supplementary Fig. 3). By using the MSCC, up to three sgRNAs could be cloned into the vector in a one-pot reaction with high efficiency. If the number of sgRNA is greater than three, pre-assembly of the inserts is recommended.

### CBE systems can induce C-to-T base conversion in *S. coelicolor*

We first tested the editing potential of two CBEs, BE2 (APOBEC1-dCas9-UGI) and BE3 (APOBEC1-nCas9-UGI) in the *Streptomyces* model strain *S. coelicolor* M145. Two target genes, *redD* and *actI-ORF2*, were selected. The gene *redD* encodes a pathway-specific activator for the biosynthesis of the red-pigmented antibiotic undecylprodigiosin (RED), and *actI-ORF2* encodes a polyketide chain length factor involved in the biosynthesis of the blue-pigmented antibiotic γ-actinorhodin (γ-ACT). sgRNAs targeting *redD* and *actI-ORF2* were designed within the first half of the ORFs to introduce premature stop codons (Fig. 2a). Before incorporating these two sgRNAs into one plasmid, we cloned each of them into separate plasmids to examine the performance of BE2 and BE3 in targeting single genes.

**Fig. 2.**
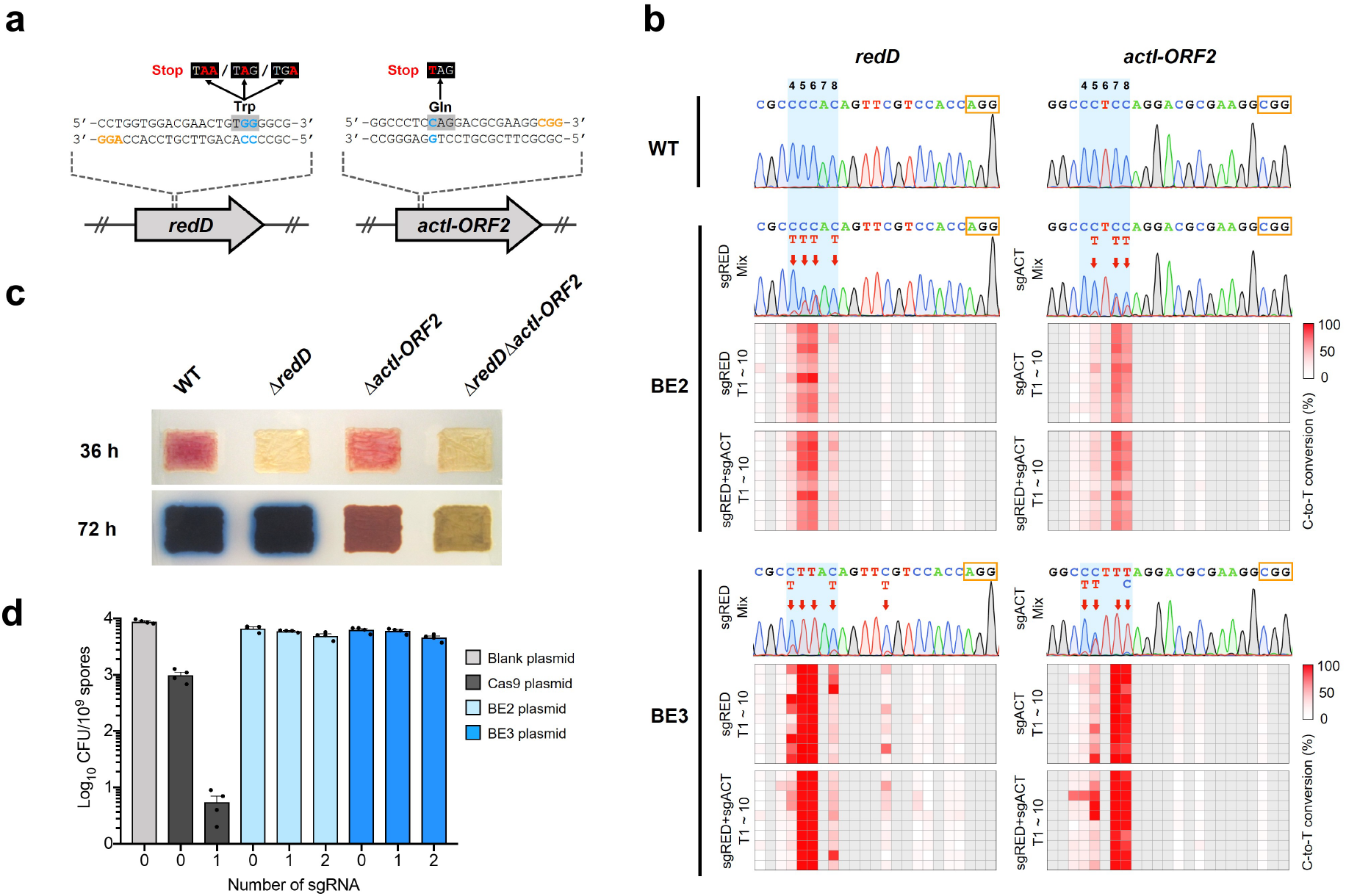
CBE-mediated genome editing in *S. coelicolor.* **a** Target design to introduce premature stop codon in *redD* and *actI-ORF2*. The NGG PAMs are shown in orange. The C•G → T•A base-pair conversions to generate premature early stop codons are highlighted in blue and red with corresponding codons shaded. **b** Base editing in *redD* and *actI-ORF2* was monitored by Sanger sequencing. Chromatograms represent WT and the editing event in the mixture of over 100 transconjugants. The NGG PAMs are framed in orange. The 4-8 nt editing window is shadowed in blue. The red arrows indicate the edited nucleotide. Experiments were repeated three times with similar results. The matrix below the chromatograms reflects the base editing events in single transconjugants. Ten transconjugants of each group were subjected to Sanger sequencing. Colour represents the percentage of total peak area with C-to-T conversion. **c** Phenotyping of representative regenerated mutants. Two pigmented antibiotics are monitored at different stages. The red-pigmented antibiotic RED could be observed after growth for 36 h on R5-medium plates, after which the blue-pigmented antibiotic γ-ACT supplants the original red colour at 72 h. **d** Survival rate in conjugal mating using the CRISPR-Cas system and CBE systems. Values and error bars reflect mean ± s.d. of four biological replicates.

To make an overall assessment of the editing efficiency in a large population, genomic DNA was extracted from a mixture of over 100 transconjugants. PCR and Sanger sequencing revealed obvious C-to-T conversions in the approximately 5-nt wide editing window (4^th^ to 8^th^ of the 20-nt protospacer region), indicating that both CBEs are functional (Fig. 2b). As judged by the relative peak areas of the sequencing chromatogram, BE3 showed higher efficiency than BE2. BE3 induced almost complete C-to-T conversions at position 5-6 for *redD* and at position 7-8 for *actI-ORF2*, while use of BE2 resulted in C/T overlapping peaks.

The editing event in single transconjugants was subsequently investigated. Surprisingly, all of the sequenced transconjugants (10/10) were found to contain edited genomes, even for the BE2-derived transconjugants (Fig. 2b and Supplementary Fig 4). The C/T overlapping peaks observed in the mixed population were also detected in the single transconjugants, so the edited and non-edited genomes apparently co-exist within a transconjugant colony. A possible explanation could be that in *Streptomyces* mycelium multiple (up to 50) genomes may share the same cytoplasmic compartment.

To demonstrate an unbiased editing efficiency and regenerate pure mutants, single colonies were isolated from the mixed transconjugants after non-selective growth. The procedure of sporulation enables the genomes of different genotypes among the mycelium to separate from each other and to be packaged into single spores. At the same time, plasmid curing takes place because the *Streptomyces* replicon (pIJ101 replicon) we used is segregationally unstable in the absence of antibiotic selection pressure^33^. Replica plating revealed that over 90% of the colonies lost apramycin resistance, showing that the plasmid can be eliminated easily after editing. Then, 30 colonies of each group were genotyped by Sanger sequencing. As expected each colony was genetically uniform and no overlapping peaks were detected at this stage. For BE3-guided mutagenesis, the C-to-T mutation was observed at a frequency of 100% (30/30) at the target site in both *redD* and *actI-ORF2* genes. In contrast, for BE2, only 43% (13/30) and 53% (16/30) colonies at the target site of *redD* and *actI-ORF2*, respectively, harbour the C-to-T mutation. A few unexpected nucleotide changes (C-to-G and C-to-A mutation) and indel mutations were found at a total frequency of 5% (3/60) resulting from BE3-guided mutagenesis (Supplementary Fig. 5).

Consistent with their genotype, in all the mutants bearing premature stop codons the production of the corresponding pigmented antibiotics was abolished (Fig. 2c and Supplementary Fig. 5b). This was confirmed by LC-ESI-HRMS analysis (Supplementary Fig. 6). These data demonstrate the applicability of two CBE systems in *S. coelicolor* as a simple and reliable way to inactivate the target gene by introducing a premature stop codon, and they also reveal that BE3 acts more efficiently than BE2 in targeting single genes.

### CBE systems permit dual-target genome editing in *S. coelicolor* with high survival rate

Next, we incorporated two sgRNAs that target *redD* and *actI-ORF2* into a single plasmid to further test the feasibility of dual-target genome editing. The conventional CRISPR/Cas system utilizes the DSB as a counter-selection marker to kill non-edited cells, and thus achieves an up to 100% editing efficiency in *Streptomyces.* However, since bacterial cells cannot readily repair DSB, efficient multiplex genome editing using the CRISPR/Cas system remains a huge challenge. We reasoned that BE systems which do not generate DSB could be used to edit the genome with less survival pressure, and would favour simultaneous multiplex genome editing. We compared the survival rate of recombinants from CRISPR/Cas and from two different CBE systems in conjugal mating experiments in *S. coelicolor*. It was observed that introduction of the no-sgRNA Cas9 plasmid caused a 10-fold decrease in the number of transconjugants compared to the blank plasmid (Fig. 2d), possibly due to the toxicity of overexpressing the exogenous DNA endonuclease. When the sgRNA was introduced and DSB was generated on the chromosome DNA, only a handful of transconjugants could be obtained by the repair system. In contrast, the no-sgRNA CBE plasmids did not significantly reduce transconjugant survival. When either one or two sgRNA were added, the survival rate only slightly decreased. Importantly, despite the nCas9-guided editor BE3 creating a SSB on the genomic DNA, its survival rate was comparable with the dCas9-guided editor BE2, indicating that one or two nicks generated during base editing are tolerated by the *Streptomyces* cell. Further, while BE2 showed an independent mutation efficiency of 70% (21/30) or 57% (17/30) and a dual-target efficiency of 43% (13/30), BE3 achieved 100% (30/30) mutation efficiency on both targets (Fig. 2b, Supplementary Figs. 4 and 5). Mutants bearing premature stop codons on both *redD* and *actI-ORF2* simultaneously abolished the production of both pigmented antibiotics (Fig. 2d and Supplementary Fig. 5). These results demonstrated that BE2 and (especially) BE3 can simultaneously edit two targets with a high survival rate and high editing efficiency.

### Multiplex genome editing by BE3 in *S. avermitilis*

Encouraged by these results, we further investigated the use of BE3 in multiplex genome editing. A proof of concept experiment was conducted in *S. avermitilis* MA-4680, the producing strain of the globally-used anthelmintic and insecticidal compound avermectin (AVE). In addition to the avermectin biosynthetic gene cluster (*ave*), there are eleven other PKS-containing clusters scattered across the *S. avermitilis* genome^34^ (Fig. 3a). We sought to demonstrate the power of this technology by simultaneously destroying all the other PKS gene clusters, hoping to block the presumptive competition for common precursors shared among different PKS pathways and hence increase the level of production of AVE.

**Fig. 3.**
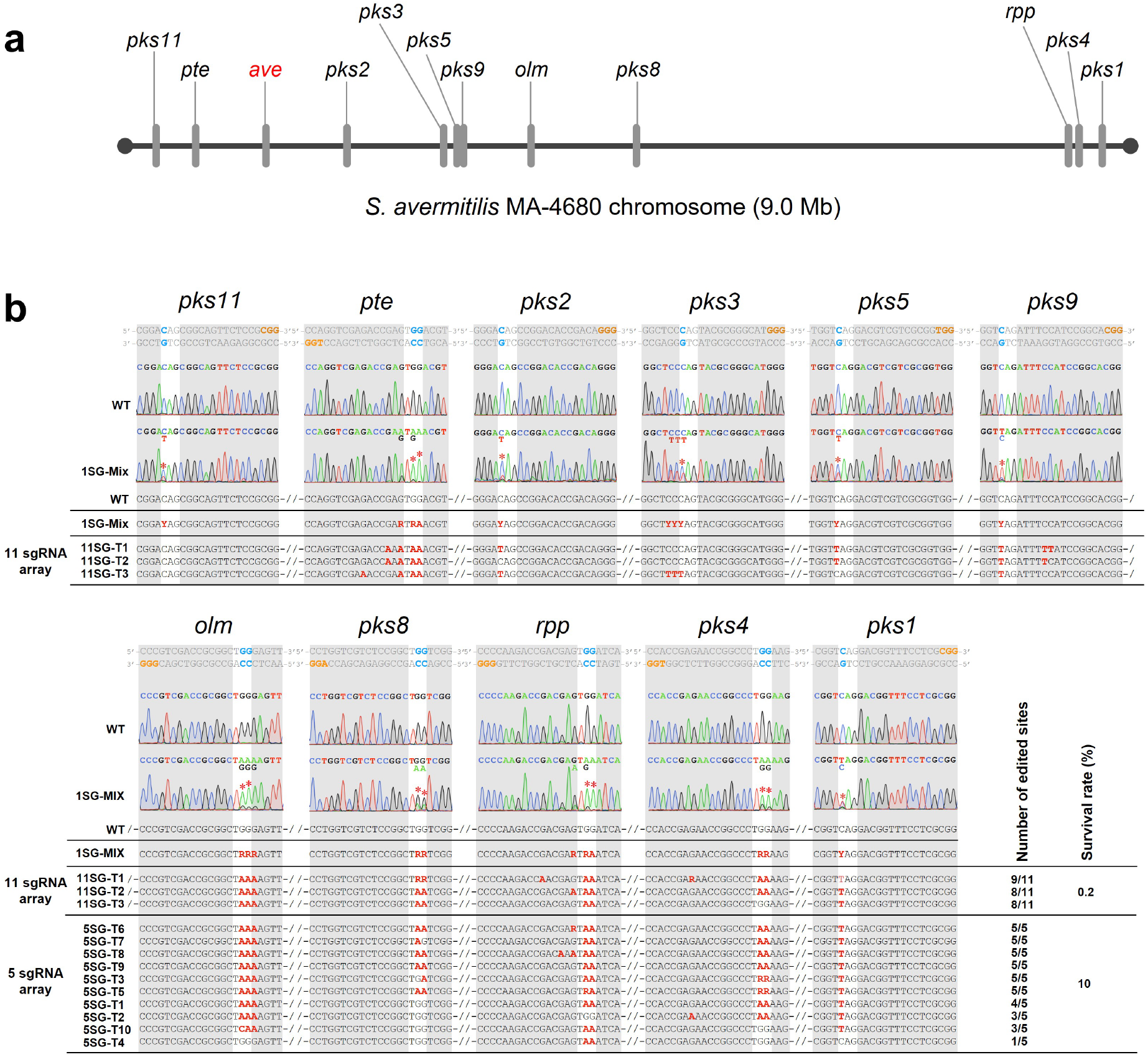
BE3 mediated multiplex genome editing in *S. avermitilis.* **a** Distribution of the twelve PKS gene clusters across the *S. avermitilis* MA-4680 chromosome. **b** Result of multiplex editing. Double strand DNA sequence on the top show the target design. The NGG PAMs and the C•G → T•A base-pair conversions to generate premature early stop codon are shown in orange and blue, respectively. Sanger sequencing chromatograms show the independent editing efficiencies of each sgRNA (1SG-Mix means the DNA sample is from the mixture of over one hundred of 1-target transconjugants). The desired base conversions to generate premature early stop codon are labeled by stars. Alignments at the bottom show the genotype of the 5-sgRNA and 11-sgRNA transconjugants.

We first designed sgRNAs to introduce premature stop codons in essential PKS genes of these 11 PKS gene clusters (3 sgRNAs were designed for each cluster) and confirmed them to be functional individually. The highest editing efficiencies were comparable to the results in *S. coelicolor*, and on these targets BE3 led to almost complete C-to-T conversion. However, lower editing efficiencies were also observed (Supplementary Fig. 7 and Supplementary Table. 2). The data allowed us to analyze the correlation between the target sequence context and the editing efficiency. It was observed that BE3 prefers the 5’-T<u>C</u> motif and discriminates against the 5’-G<u>C</u> motif (the target C is underlined) (Supplementary Fig. 8a). This sequence preference of BE3 has been established in previous work^10^, and the mechanism has been explained by the protein structure of the APOBEC family deaminase^35^. The most efficient editing occurred at position 6 in the 4-8^th^ window and a gradually decreased efficiency was observed towards the flanking positions (Supplementary Fig. 8b). Importantly, using 20-nt spacers with a G+C content higher than 75% (number of G+C>15) resulted in a lower editing efficiency (Supplementary Fig. 8c).

Plasmids comprising the most effective sgRNAs for each of either five or eleven target gene clusters were constructed and delivered into *S. avermitilis* (Fig. 3b). The result showed that six out of the ten sequenced 5-target transconjugants had been edited in all five target sites, while between one and four targets were edited in the remaining four transconjugants, giving a 5-target simultaneous mutation rate of 60%. Moreover, up to eight or nine targets were found to be edited simultaneously in the three 11-target transconjugants that were obtained. For comparison, the survival rate using a 2-sgRNA plasmid was 73% in *S. coelicolor*, while the 5-sgRNA and 11-sgRNA plasmids resulted in sharp decrease in *S. avermitilis* survival rate (10% for the 5-sgRNA plasmid and 0.2% for the 11-sgRNA plasmid).

The production of AVE in the transconjugants was determined. Three 5-target mutants (5SG-T6, 5SG-T7, 5SG-T8) and three 11-target mutants (11SG-T1, 11SG-T2 and 11SG-T3) were subjected to fermentation and LC-ESI-HRMS analysis. However, only 5SG-T7 showed an insignificant increase in AVE production (Supplementary Fig. 9) while in the rest of the mutants production of AVE was abolished. This may reflect off-target editing in the *ave*, particulary in the highly homologous PKS genes. Using the DNA sample of 1-sgRNA transconjugant mixture, we investigated the eight most likely off-target sites (OTs) in the *ave* (Supplementary Table. 3), and found that only sgRNA targeting the oligomycin biosynthetic gene cluster (*olm*) induced detectable off-target editing on three OTs (*olm*-OT-4, *olm*-OT-5, *olm*-OT-6) (Supplementary Fig.10). These three OTs, bearing four nucleotide mismatches compairing with the on-target site, have identical sequence. The off-target editing here would generate stop codons at *aveA4* gene and abolish the avermectin synthesis (Fig. 4a). Then, we investigated the mutants and proved the relevance between the off-target editing and the AVE abolishment (Supplementary Fig.10). Overall, these results exhibited that the BE3 enable a highly efficient multiplex mutagenesis in *Streptomyces*, and enough attention should also be given to the off-target effect in targeting promiscuous sites with highly homology.

**Fig. 4.**
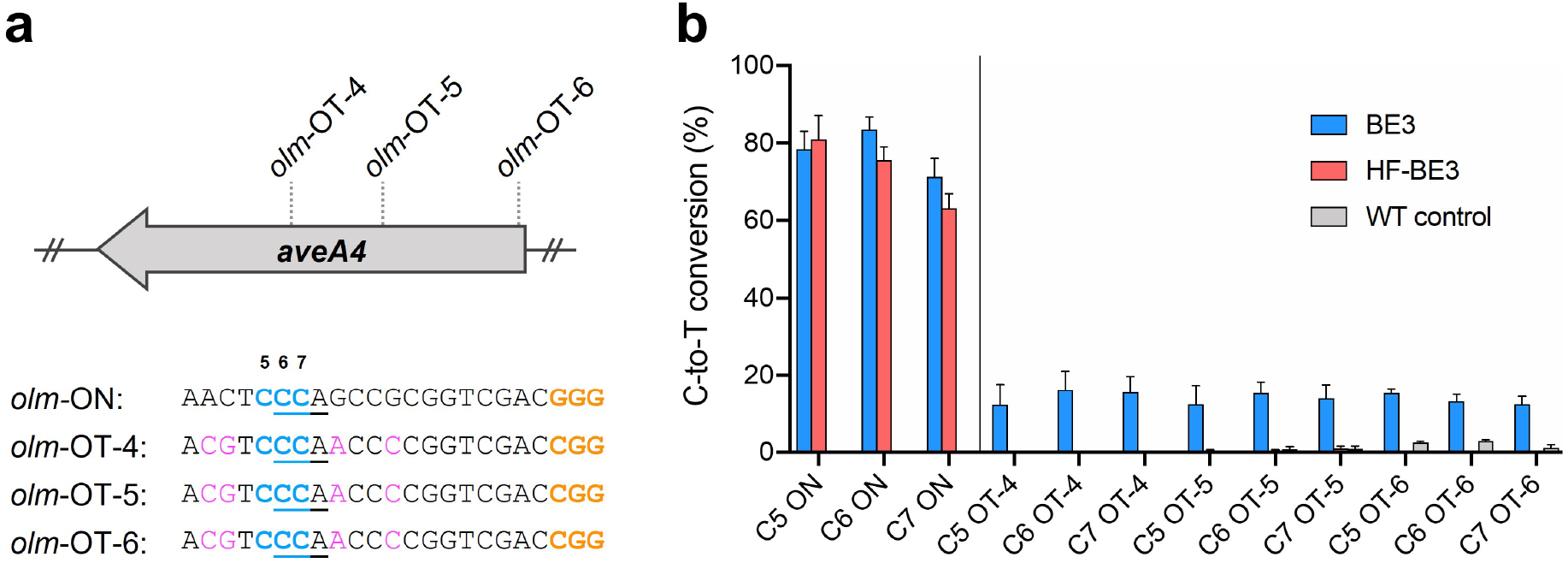
On-and off-target activity of BE3 and HF-BE3 **a** Location and sequence of the three OTs associated with sgOLM-1 at *aveA4* gene. PAM, mismatched bases and target Cs in editing window are shown in orange, magenta and blue, respectively. Codons to generate premature stop codons are underlined. **b** Editing efficiency of BE3 and HF-BE3 was assessed by Sanger sequencing. On- and off-target loci are separated by a vertical line. Values and error bars reflect mean ± s.e.m.; n=3. For each biological repeat, genomic DNA was extracted from the mixture of over 100 tranconjugants. Minimal conversions in the WT control are attributed to the background noise of Sanger sequencing.

### Improved editing specificity by using HF-BE3

Although off-target editing could be avoided in most cases by careful design of sgRNA, the application of BE3 to very challenging sites, e.g. the highly repetitive modular PKS genes, is inevitably limited. High-fidelity BE3 (HF-BE3)^13^, in which the original nCas9 was replaced by a high-fidelity nCas9 variant (HF-nCas9)^36^, has been reported to reduce the off-target effect by decreasing non-specific DNA contacts. Therefore, we generated a version of HF-BE3 by introducing the four HF mutations (N497A, R661A, Q695A and Q926A) to the *Streptomyces* codon-optimized BE3, and repeated the *olm* disruption assay with the sgRNA used in the previous experiment. Assessment of the C-to-T conversion efficiency by Sanger sequencing revealed comparable on-target activities with BE3 and HF-BE3. Strikingly, installation of the HF mutations reduced the off-target editing at *olm*-OT-4, *olm*-OT-5, *olm*-OT-6 from an average 14% to an undetectable level (Fig. 4b and Supplementary Fig. 10).

### The ABE system can induce A-to-G base conversion in *S. coelicolor*

The ABE system which enables targeted A•T → G•C base-pair conversion expands the choice of amino acid conversions. To explore the editing potential of the ABE system in *Streptomyces*, we tested the well-established editor ABE7.10^28^ in both the dCas9 and nCas9 versions (referred to as ABEd and ABEn in this study, Fig. 1b) in *S. coelicolor.* The target gene, *actVB*, encodes a flavin reductase that is involved in the dimerization of two 6-deoxy-dihydrokalafungin (DDHK) molecules to form γ-ACT, so deletion of *actVB* leads to accumulation of the brown-pigmented shunt product actinoperylone (ACPL)^37^. According to the approximately 4-nt wide editing window (4^th^ to 7^th^ of the 20-nt protospacer region) of ABE7.10^28^, the base-pair conversion here was designed to mutate its start codon (ATG) to a threonine codon (ACG) thus disrupting the initiation of *actVB* translation (Fig. 5a). After conjugal transfer, the expected A-to-G conversion was observed by Sanger sequencing, indicating the functionality of the ABE system (Fig. 5b). Consistent with the above observations in CBE, the editing efficiency displayed a difference between dCas9- and nCas9-guided ABEs. While the mixture of over 100 transconjugants and ten independently investigated transconjugants showed the A/G overlapping peak by using ABEd, complete A-to-G conversion was achieved in ABEn derived transconjugants to give a mutation efficiency of 100% (Fig. 5b and Supplementary Fig. 11). As we expected, both visual observation and LC-ESI-HRMS analysis confirmed the accumulation of ACPL in the mutant (Fig. 5c). In addition, no reduced survival rate in conjugal transfer was observed for the ABE plasmids, suggesting their potential for multiplexing. These results indicate that the ABE system is efficient in *Streptomyces*.

**Fig. 5.**
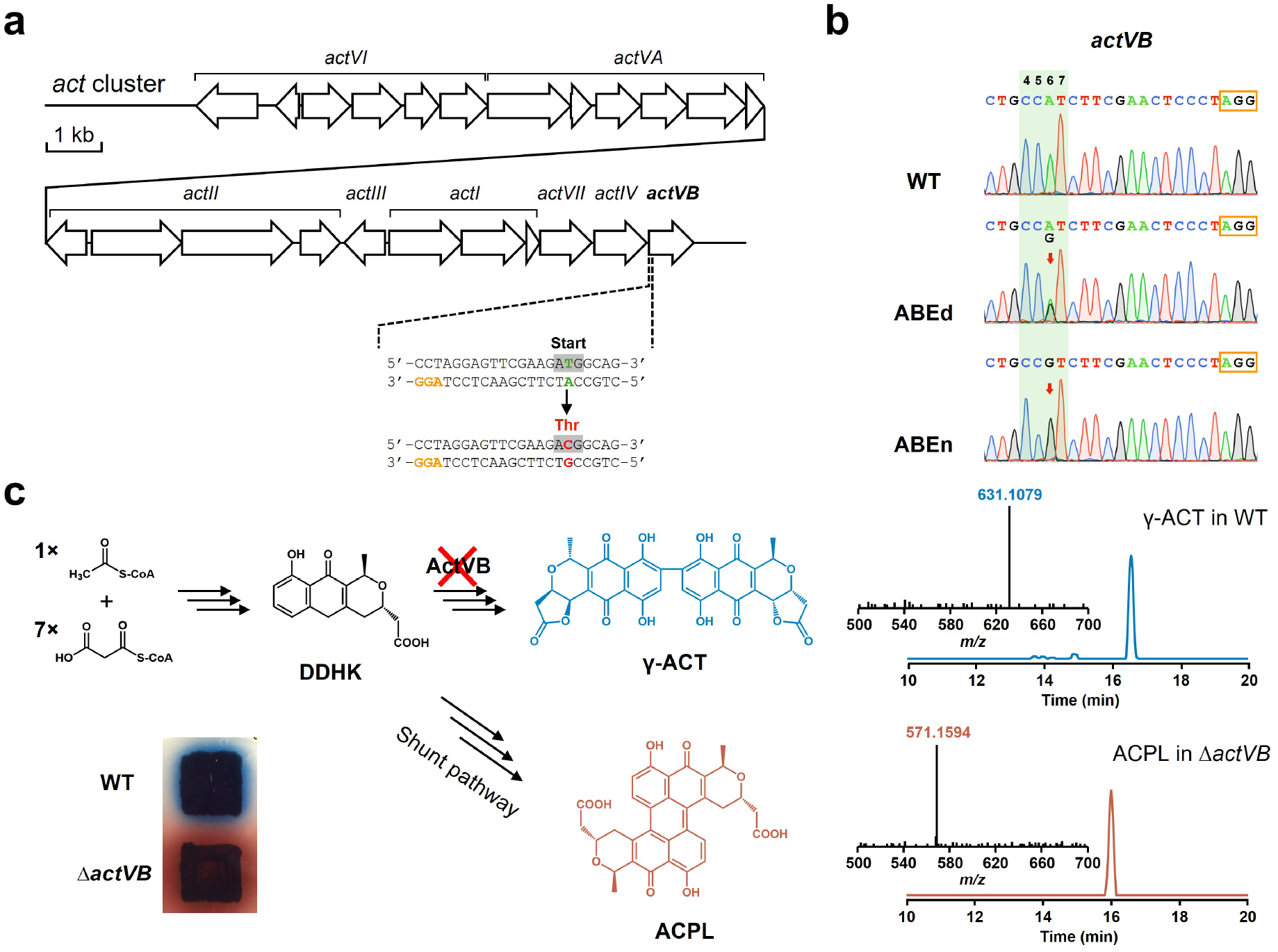
ABE mediated genome editing in *S. coelicolor.* **a** Target design to mutate the strat codon of *actVB*. The NGG PAM is shown in orange. Targeted A•T → G•C base-pair conversion to mutate the strat codon is highlighted in green and red with corresponding codons to be shaded. **b** Base editing at *actVB* was assessed by Sanger sequencing. Chromatograms represent WT and the editing event in the mixture of 100 transconjugants. The NGG PAMs are framed in orange. The 4-7 nt editing window is shadowed in green. The red arrows indicate the edited nucleotide. Experiments were repeated three times with similar results. Editing events in single transconjugants are supplied in Supplementary Fig. 11. **c** Detection of the brown pigmented shunt product actinoperylone (ACPL) by visual observation and LC-ESI-HRMS.

## Discussion

In this work, we constructed a series of *Streptomyces* base editing plasmids that allowed for convenient single or multiple sgRNA cloning. Both CBEs and ABEs achieved successful editing in *Streptomyces*, with an up to 100% editing efficiency by using the nCas9-guided editors compared to approximately 50% efficiency by using the dCas9-guided editors. In comparison to the CRISPR/Cas system and to time-consuming mutant screening by traditional homologous recombination, the high survival rate in conjugal mating suggests a huge advantage of using the BE systems for genome editing in *Streptomyces* species. Consistent to part of our study, the BE3 was also investigated to serve as programmable base editor in *Streptomyces* by Tong et al. during our manuscript preparation^38^.

We further investigated the feseability and efficiency of BE3 in synchronous multiplex genome editing. A 5-target editing efficiency of 60% was acheived, and editing of up to nine targets in the 11-target mutagenesis was obtained. The increased target number inevitably resulted in fewer surviving transconjugants, but to our surprise the editing efficiency was not affected. We found that the editing efficiency of BE3 depends on the sequence context of the target sites, in addition to the 5’-motif discrimination of APOBEC1, the extremely high G+C content of the target site also affects editing efficiency. This observation is especially informative for target selection in the high G+C *Streptomyces*. Since undesired off-target editing was detected by using promiscuous sgRNA, HF-BE3 can be used to circumvent it effectively.

When testing the aility of the ABE systems in *Streptomyces*, we targeted a start codon and successfuly abolished the function of the corresponding gene, thus providing an alternative strategy for gene disruption. However, compared to the method of introducing premature stop codon to the full length ORF, mutating a start codon is often limited by the requirement of a nearby NGG PAM. We look forward the use of a Cas9 variant that recognizes the NG PAM (SpCas9-NG)^39^ will expand the editable sites and make the strategy universally applicable. In addition, translation of many *Streptomyces* secondary metabolite genes is profoundly restricted by the rare leucine codon TTA^40^, this codon can also be optimized by ABE.

It is also worth noting that although the dCas9-guided base editors only have half the efficiency of the nCas9 versions, they are convenient to generate single- and multiple-target mutants if the guide sequences can be appropriately designed.

In summary, we have demonstrated that the CRISPR/Cas9-deaminase base editors provide a rapid and effective method for genetic manipulation in *Streptomyces*. Its adaptation for use in this genus is expected to accelerate functional genetic studies of these species, thus further promoting the discovery and engineering of valuable natural products.

## Methods

### Bacterial strains, culture conditions and reagents

*E. coli* DH10B was used for plasmid construction. *E. coli* ET12567 harboring pUZ8002 was used as the donor for intergeneric conjugation between *E. coli* and *Streptomyces*. *E. coli* strains were maintained in 2×YT medium^2^ at 37°C, antibiotics (apramycin 25 μg/mL, ampicillin 100 μg/mL, chloramphenicol 25 μg/mL, and kanamycin 25 μg/mL) were added into the medium when required. *Streptomyces* strains were cultivated on SFM agar medium^2^ at 28°C for growth of mycelium and sporulation. SFM and ISP4 agar medium^41^ with addition of 10 mM MgCl_2_ were used for the conjugation of *S. coelicolor* M145 and *S. avermitilis* MA-4680, respectively. R5 agar medium^2^ was used for RED, γ-ACT and ACPL production. TSBY liquid medium^2^ and avermectin fermentation liquid medium^42^ were used for avermectin production. Plasmids and DNA oligos used in this study are listed in Supplementary Tables 4 and 5.

### Plasmids construction

A *Streptomyces*-codon-optimized *cas9* gene was synthesized in our previous study^6^, DNAs encoding the remaining components (APOBEC1-Cas9_N_terminal-UGI for CBE; TadA-TadA*-Cas9_N_terminal for ABE) to complete the base editors were codon-optimized for *Streptomyces*, then together with the *rpsL*p and *fd*t the designed sequences were synthesized by GenScript and provided on the vector pUC57 to create transitional plasmids pUC57-SCBE-wocas9 and pUC57-SABE-wocas9, respectively. DNAs encoding the remaining portion of the nCas9 or dCas9 were PCR amplified from pWHU2650^6^ and inserted into the above transitional plasmids by Gibson assembly. The common D10A mutation of nCas9 and dCas9 was incorporated in the synthesized Cas9 N-terminal; the H840A mutation of dCas9 was introduced by an additional overlap PCR. The multiple sgRNA cloning cassette (MSCC, *Xba*I-sgRNA_scaffold-B1006t-*kasO*p*-*Nhe*I) was also synthesized and provided on pUC57. The MSCC was cloned into pYH7-wocos, a shortened derivative of the *E. coli*-*Streptomyces* shuttle vector pYH7^33^, in which a 1.2 kb fragment including the *cos* site was removed, to give the plasmid pYH7-MSCC. Four versions of base editors were incorporated into pYH7-MSCC to give the *Streptomyces* multiplex base editing plasmids pSCBE2, pSCBE3, pSABEd and pSABEn. Based on the MSCC containing base editing plasmids, the single sgRNA cloning cassette (SSCC, *kasO*p*-*Bae*I-B1006t) was constructed by Gibson assembly to give the plasmids pSCBE2-single, pSCBE3-single, pSABEd-single and pSABEn-single that more suitable for single sgRNA cloning. The HF mutations were introduced to pSCBE3 and pSCBE3-single by *in vitro* Cas9 digestion^43^, overlap PCR and Gibson assembly, creating pSCBE3-HF and pSCBE3-HF-single.

### sgRNA design and cloning

The sgRNAs for CBE were designed using the Benching web tool (https://benchling.com/), in which the criterion for scoring base editing guide sequence was incorporated according to the *in vitro* activity of BE1. The sgRNA for ABE was designed manually. The off-target risk of the candidate sgRNAs was evaluated by CasOT^44^, sgRNAs with higher specificity were selected. Because most sgRNAs used in this study needed to be cloned individually and connected in series, the MSCC was used for all sgRNA cloning to share primers. Cloning steps were as follows: (1) linearize the target BE plasmid with *Xba*I and *Nhe*I. (2) PCR amplify the inserts using the chosen MSCC containing BE plasmids as template, and introduce the 20 bp guide sequence and the 20 bp vector overlaps by the 5’ end of the primers. (3) use Gibson assembly to introduce the inserts into the *Xba*I-*Nhe*I linearized BE plasmid to make functional sgRNA expression cassettes, then check the recombinants by colony PCR and Sanger sequencing.

### Conjugation delivery of plasmids into *Streptomyces*

Conjugation experiments were undertaken following the standard protocol described previously^2^. Density of the spore suspension was determined by the hemacytometer counting method. The number of transconjugants was counted after growth on the conjugation plate for seven days.

### Editing efficiency evaluation

Genomic DNA was extracted from the mycelium at appropriate stages. To evaluate the editing efficiency in a large base population, the genomic DNA was directly extracted from the mixture of over 100 transconjugants scraped out from the conjugation plates after overlaying with apramycin for 7 days. To examine the editing event in single transconjugants, the randomly selected transconjugants were given an extra 3 days growth on the SFM agar without apramycin to produce enough genomic DNA. To genotype the regenerated plasmid-free mutants, the mixture of over 100 transconjugants was inoculated onto SFM agar without apramycin and grow for 7 days for sporulation. The spores were then harvested and spread onto SFM agar without apramycin in serial dilution to achieve single colony. The resulting single colonies were replica plated onto the plates with and without apramycin for phenotype screening. Genomic DNA was extracted from the strains without apramycin resistance. For all genomic DNA samples, target sites were PCR amplified with Phusion High-Fidelity DNA Polymerase (ThermoFisher), then subjected to Sanger sequencing by GenScript and analyzed using SnapGene viewer (https://www.snapgene.com/). The percentage of base conversion was estimated by EditR, a software to quantify the base editing efficiency from Sanger sequencing chromatograms^45^.

### Production, isolation, and analysis of secondary metabolites

For the production of pigmented antibiotics (RED, γ-ACT, ACPL), *S. coelicolor* and its mutants were plated onto R5 agar medium and incubated at 28°C. The photograph was taken from the back of the plate at 36 and 72 hours. For the fermentation of AVE, *S. avermitilis* and its mutants were cultured in two stages. The seed culture was maintained in TSBY liquid medium at 28°C with shaking at 220 rpm for 2 days, then inoculate 10% seed into the fermentation liquid medium and incubated at 28°C with shaking at 220 rpm for 7 days. The plate culture of *S. coelicolor* and the liquid cultre of *S. avermitilis* were treated overnight with an equal volume of acidified methanol (0.1% formic acid) or three volumes of methanol, respectively. The supernatants of the extracts were then subjected to liquid chromatography-electrospray ionization-high-resolution mass spectrometry (LC-ESI-HRMS) analysis on an LTQ Orbitrap XL (Thermo Fisher) using positive-mode electrospray ionization. Analysis was performed with a Phenomenex Luna C18 column (5 μm, 250×4.6 mm) at a flow rate of 1 mL/min using a mobile phase of (A) 0.1% trifluoroacetic acid in water and (B) 0.1% trifluoroacetic acid in acetonitrile. The gradient for separation of RED, γ-ACT, ACPL: 0–2 min 5% B, 2–17 min 5% B to 100 % B, 17–22 min 100% B, 22–23 min 100% B to 5% B, 23–27 min 5% B. For detection of AVE components, flow rate is 0.6 mL/min using a mobile phase of 1(A) 0.1% formic acid in water and (B) acetonitrile. The gradient for separation of AVE components: 0–27 min 88% B, 27–27.5 min 88% B to 10% B, 27.5–29 min 10% B, 29–29.5 min 10% B to 98% B, 29.5–31 min 98% B, 31–31.5 min 98% B to 88% B, 31.5–36 min 88% B.

## Data availability

All the data generated or analyzed during this study are included in this published article.

## Acknowledgements

This work was supported by the National Natural Science Foundation of China (31770069). The authors thank Prof. Peter F. Leadlay at University of Cambridge for his critical reading of the manuscript.

## Author contributions

Y.S. and Z.Z. conceived the idea and designed the experiments; Z.Z., J.G., L.D., L.C., J.W. and S.L. conducted the experiments; All the authors analyzed the results; Z.Z. and Y.S. wrote the manuscript.

## Author information

These authors contributed equally: Zhiyu Zhong and Junhong Guo.

## References

1. Hopwood, D. A. Streptomyces in Nature and Medicine: The Antibiotic Makers. (Oxford University Press, New York, 2007).

2. Kieser, T. et al. Practical Streptomyces Genetics. (John Innes Foundation, Norwich, 2000).

3. Cobb, R. E., Wang, Y. & Zhao, H. High-efficiency multiplex genome editing of *Streptomyces* species using an engineered CRISPR/Cas system. ACS Synth. Biol. 4, 723–728 (2015).

4. Huang, H., Zheng, G., Jiang, W., Hu, H. & Lu, Y. One-step high-efficiency CRISPR/Cas9-mediated genome editing in *Streptomyces*. Acta Biochim. Biophys. Sin. 47, 231–243 (2015).

5. Tong, Y., Charusanti, P., Zhang, L., Weber, T. & Lee, S. Y. CRISPR-Cas9 based engineering of actinomycetal genomes. ACS Synth. Biol. 4, 1020–1029 (2015).

6. Zeng, H. et al. Highly efficient editing of the actinorhodin polyketide chain length factor gene in *Streptomyces coelicolor* M145 using CRISPR/Cas9-CodA(sm) combined system. Appl. Microbiol. Biotechnol. 99, 10575–10585 (2015).

7. Zhang, M. M. et al. CRISPR-Cas9 strategy for activation of silent *Streptomyces* biosynthetic gene clusters. Nat. Chem. Biol. 13, 607–609 (2017).

8. Li, L. et al. CRISPR-Cpf1 assisted multiplex genome editing and transcriptional repression in *Streptomyces*. Appl. Env. Microbiol. 84, e00827–18 (2018).

9. Cui, L. & Bikard, D. Consequences of Cas9 cleavage in the chromosome of *Escherichia coli*. Nucleic Acids Res. 44, 4243–4251 (2016).

10. Komor, A. C., Kim, Y. B., Packer, M. S., Zuris, J. A. & Liu, D. R. Programmable editing of a target base in genomic DNA without double-stranded DNA cleavage. Nature 533, 420–424 (2016).

11. Nishida, K. et al. Targeted nucleotide editing using hybrid prokaryotic and vertebrate adaptive immune systems. Science 353, aaf8729 (2016).

12. Kim, Y. B. et al. Increasing the genome-targeting scope and precision of base editing with engineered Cas9-cytidine deaminase fusions. Nat. Biotechnol. 35, 371–376 (2017).

13. Rees, H. A. et al. Improving the DNA specificity and applicability of base editing through protein engineering and protein delivery. Nat. Commun. 8, 15790 (2017).

14. Komor, A. C. et al. Improved base excision repair inhibition and bacteriophage Mu Gam protein yields C:G-to-T:A base editors with higher efficiency and product purity. Sci. Adv. 3, eaao4774 (2017).

15. Li, X. et al. Base editing with a Cpf1–cytidine deaminase fusion. Nat. Biotechnol. 36, 324–327 (2018).

16. Gehrke, J. M. et al. An APOBEC3A-Cas9 base editor with minimized bystander and off-target activities. Nat. Biotechnol. 36, 977–982 (2018).

17. Zhang, Y. et al. Programmable base editing of zebrafish genome using a modified CRISPR-Cas9 system. Nat. Commun. 8, 118 (2017).

18. Kim, K. et al. Highly efficient RNA-guided base editing in mouse embryos. Nat. Biotechnol. 35, 435–437 (2017).

19. Liu, Z. et al. Highly efficient RNA-guided base editing in rabbit. Nat. Commun. 9, 2717 (2018).

20. Zong, Y. et al. Precise base editing in rice, wheat and maize with a Cas9-cytidine deaminase fusion. Nat. Biotechnol. 35, 438–440 (2017).

21. Shimatani, Z. et al. Targeted base editing in rice and tomato using a CRISPR-Cas9 cytidine deaminase fusion. Nat. Biotechnol. 35, 441–443 (2017).

22. Gu, T. et al. Highly efficient base editing in *Staphylococcus aureus* using an engineered CRISPR RNA-guided cytidine deaminase. Chem. Sci. 9, 3248–3253 (2018).

23. Tang, W. & Liu, D. R. Rewritable multi-event analog recording in bacterial and mammalian cells. Science 360, eaap8992 (2018).

24. Zheng, K. et al. Highly efficient base editing in bacteria using a Cas9-cytidine deaminase fusion. Commun. Biol. 1, 32 (2018).

25. Wang, Y. et al. MACBETH: multiplex automated *Corynebacterium glutamicum* base editing method. Metab. Eng. 47, 200–210 (2018).

26. Chen, W. et al. CRISPR/Cas9-based genome editing in *Pseudomonas aeruginosa* and cytidine deaminase-mediated base editing in *Pseudomonas* species. iScience 6, 222–231 (2018).

27. Wang, Y. et al. Precise and efficient genome editing in *Klebsiella pneumoniae* using CRISPR-Cas9 and CRISPR-assisted cytidine deaminase. Appl. Env. Microbiol. 84, e01834–18 (2018).

28. Gaudelli, N. M. et al. Programmable base editing of A•T to G•C in genomic DNA without DNA cleavage. Nature 551, 464–471 (2017).

29. Hua, K., Tao, X., Yuan, F., Wang, D. & Zhu, J.-K. Precise A·T to G·C base editing in the rice genome. Mol. Plant 11, 627–630 (2018).

30. Yan, F. et al. Highly efficient A·T to G·C base editing by Cas9n-guided tRNA adenosine deaminase in rice. Mol. Plant 11, 631–634 (2018).

31. Ryu, S.-M. et al. Adenine base editing in mouse embryos and an adult mouse model of Duchenne muscular dystrophy. Nat. Biotechnol. 36, 536–539 (2018).

32. Li, C. et al. Expanded base editing in rice and wheat using a Cas9-adenosine deaminase fusion. Genome Biol. 19, 59 (2018).

33. Sun, Y., He, X., Liang, J., Zhou, X. & Deng, Z. Analysis of functions in plasmid pHZ1358 influencing its genetic and structural stability in *Streptomyces lividans* 1326. Appl. Microbiol. Biotechnol. 82, 303–310 (2009).

34. Ikeda, H. et al. Complete genome sequence and comparative analysis of the industrial microorganism *Streptomyces avermitilis*. Nat. Biotechnol. 21, 526–531 (2003).

35. Shi, K. et al. Structural basis for targeted DNA cytosine deamination and mutagenesis by APOBEC3A and APOBEC3B. Nat. Struct. Mol. Biol. 24, 131–139 (2017).

36. Kleinstiver, B. P. et al. High-fidelity CRISPR-Cas9 variants with undetectable genome-wide off-targets. Nature 529, 490–495 (2016).

37. Okamoto, S., Taguchi, T., Ochi, K. & Ichinose, K. Biosynthesis of actinorhodin and related antibiotics: discovery of alternative routes for quinone formation encoded in the *act* gene cluster. Chem. Biol. 16, 226–236 (2009).

38. Tong, Y. et al. CRISPR-BEST: a highly efficient DSB-free base editor for filamentous actinomycetes. Preprint at bioRxiv: https://www.biorxiv.org/content/10.1101/582403v1 (2019).

39. Nishimasu, H. et al. Engineered CRISPR-Cas9 nuclease with expanded targeting space. Science 361, 1259–1262 (2018).

40. Chater, K. F. & Chandra, G. The use of the rare UUA codon to define “Expression Space” for genes involved in secondary metabolism, development and environmental adaptation in *Streptomyces*. J. Microbiol. 46, 1–11 (2008).

41. Shirling, E. B. & Gottlieb, D. Methods for characterization of *Streptomyces* species1. Int. J. Syst. Evol. Microbiol. 16, 313–340 (1966).

42. Wang, J.-B. et al. Characterization of AvaR1, an autoregulator receptor that negatively controls avermectins production in a high avermectin-producing strain. Biotechnol. Lett. 36, 813–819 (2014).

43. Liu, Y. et al. In vitro CRISPR/Cas9 system for efficient targeted DNA editing. mBio 6, e01714–15 (2015).

44. Xiao, A. et al. CasOT: a genome-wide Cas9/gRNA off-target searching tool. Bioinformatics 30, 1180–1182 (2014).

45. Kluesner, M. G. et al. EditR: a method to quantify base editing from Sanger sequencing. CRISPR J. 1, 239–250 (2018).

