## Supplementary Information for "Base editing in *Streptomyces* with Cas9-deaminase fusions"

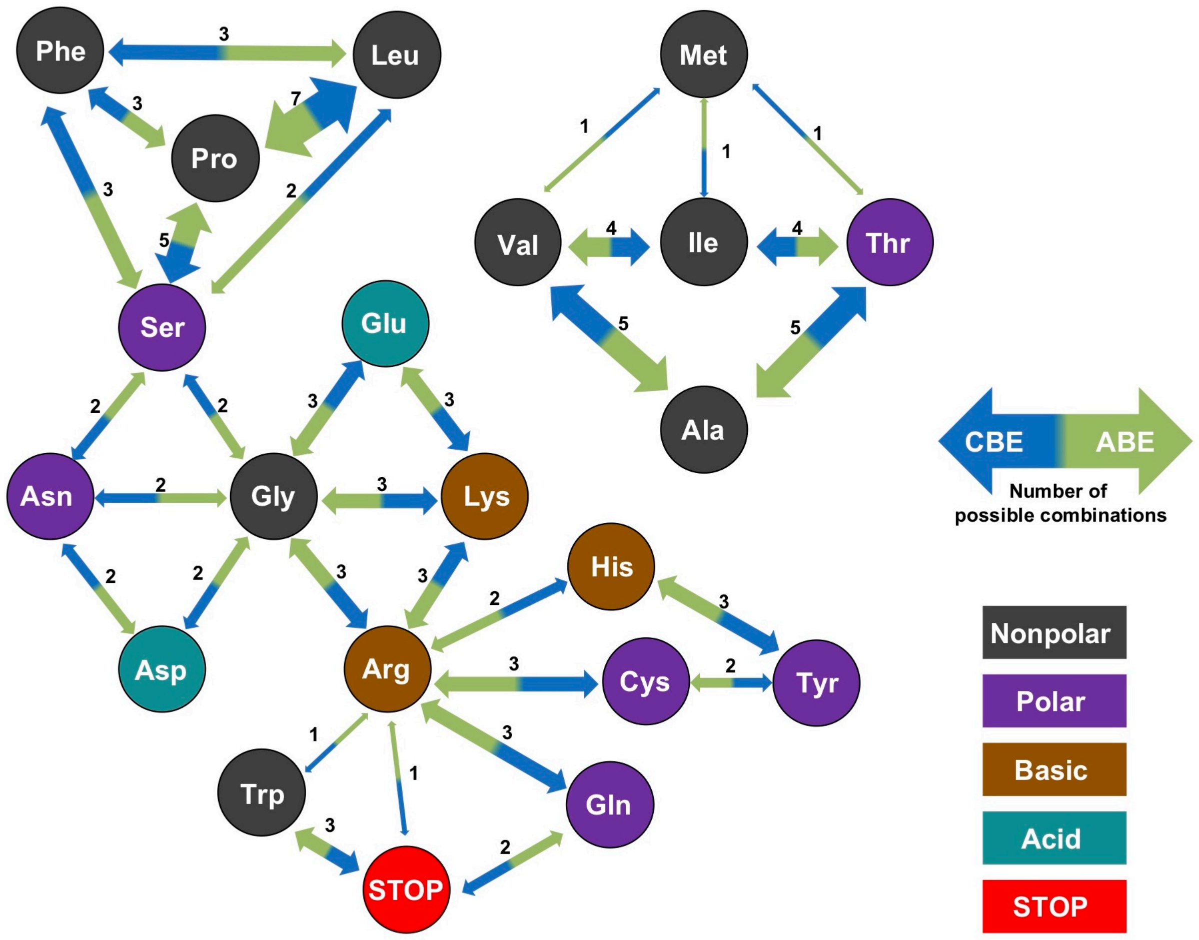


**Supplementary Fig. 1** Repertoire of the amino acid substitutions generated by CBE and ABE. Convertible amino acids are linked by two headed arrows. Blue and green arrows show the direction of amino acid substitutions generated by CBE and ABE, respectively. The number of possible combinations to generate the indicated substitutions are labeled on the arrows. Ala, alanine; Arg, arginine; Asn, asparagine; Asp, aspartic acid; Cys, cysteine; Gln, glutamine; Glu, glutamic acid; Gly, glycine; His, histidine; Ile, isoleucine; Met, methionine; Leu, leucine; Lys, lysine; Phe, phenylalanine; Pro, proline; Ser, serine; Thr, threonine; Trp, tryptophan; Tyr, tyrosine; Val, valine; STOP, STOP codon. (Figure adapted from Billon et. al 2017)

**Supplementary Table 1** Base editing codon table.

### Supplementary Table 1a. C-to-T editing

| 1st | 2nd | | | | | | | | | | | | | | | | 3rd |
|---|---|---|---|---|---|---|---|---|---|---|---|---|---|---|---|---|---|
| | T | | | | C | | | | A | | | | G | | | | |
| T | TTT | Phe | | | TCT | Ser | TTT | Phe | TAT | Tyr | | | TGT | Cys | | | T |
| | TTC | Phe | TTT | Phe | TCC | Ser | TTC, TCT, TTT | Phe, Ser, Phe | TAC | Tyr | TAT | Tyr | TGC | Cys | TGT | Cys | C |
| | TTA | Leu | | | TCA | Ser | TTA | Leu | TAA | STOP | | | TGA | STOP | | | A |
| | TTG | Leu | | | TCG | Ser | TTG | Leu | TAG | STOP | | | TGG | Trp | | | G |
| C | CTT | Leu | TTT | Phe | CCT | Pro | TCT, CTT, TTT | Ser, Leu, Phe | CAT | His | TAT | Tyr | CGT | Arg | TGT | Cys | T |
| | CTC | Leu | TTC, CTT, TTT | Phe, Leu, Phe | CCC | Pro | TCC, CTC, CCT, TTC, TCT, CTT, TTT | Ser, Leu, Pro, Phe, Ser, Leu, Phe | CAC | His | TAC, CAT, TAT | Tyr, His, Tyr | CGC | Arg | TGC, CGT, TGT | Cys, Arg, Cys | C |
| | CTA | Leu | TTA | Leu | CCA | Pro | TCA, CTA, TTA | Ser, Leu, Leu | CAA | Gln | TAA | STOP | CGA | Arg | TGA | STOP | A |
| | CTG | Leu | TTG | Leu | CCG | Pro | TCG, CTG, TTG | Ser, Leu, Leu | CAG | Gln | TAG | STOP | CGG | Arg | TGG | Trp | G |
| A | ATT | Ile | | | ACT | Thr | ATT | Ile | AAT | Asn | | | AGT | Ser | | | T |
| | ATC | Ile | ATT | Ile | ACC | Thr | ATC, ACT, ATT | Ile, Thr, Ile | AAC | Asn | AAT | Asn | AGC | Ser | AGT | Ser | C |
| | ATA | Ile | | | ACA | Thr | ATA | Ile | AAA | Lys | | | AGA | Arg | | | A |
| | ATG | Met | | | ACG | Thr | ATG | Met | AAG | Lys | | | AGG | Arg | | | G |
| G | GTT | Val | | | GCT | Ala | GTT | Val | GAT | Asp | | | GGT | Gly | | | T |
| | GTC | Val | GTT | Val | GCC | Ala | GTC, GCT, GTT | Val, Ala, Val | GAC | Asp | GAT | Asp | GGC | Gly | GGT | Gly | C |
| | GTA | Val | | | GCA | Ala | GTA | Val | GAA | Glu | | | GGA | Gly | | | A |
| | GTG | Val | | | GCG | Ala | GTG | Val | GAG | Glu | | | GGG | Gly | | | G |
| | From | | To | | From | | To | | From | | To | | From | | To | | |

### Supplementary Table 1b. G-to-A editing

| 1st | 2nd | | | | | | | | | | | | | | | | 3rd |
|---|---|---|---|---|---|---|---|---|---|---|---|---|---|---|---|---|---|
| | T | | | | C | | | | A | | | | G | | | | |
| T | TTT | Phe | | | TCT | Ser | | | TAT | Tyr | | | TGT | Cys | TAT | Tyr | T |
| | TTC | Phe | | | TCC | Ser | | | TAC | Tyr | | | TGC | Cys | TAC | Tyr | C |
| | TTA | Leu | | | TCA | Ser | | | TAA | STOP | | | TGA | STOP | TAA | STOP | A |
| | TTG | Leu | TTA | Leu | TCG | Ser | TCA | Ser | TAG | STOP | TAA | STOP | TGG | Trp | TAG, TGA, TAA | STOP, STOP, STOP | G |
| C | CTT | Leu | | | CCT | Pro | | | CAT | His | | | CGT | Arg | CAT | His | T |
| | CTC | Leu | | | CCC | Pro | | | CAC | His | | | CGC | Arg | CAC | His | C |
| | CTA | Leu | | | CCA | Pro | | | CAA | Gln | | | CGA | Arg | CAA | Gln | A |
| | CTG | Leu | CTA | Leu | CCG | Pro | CCA | Pro | CAG | Gln | CAA | Gln | CGG | Arg | CAG, CGA, CAA | Gln, Arg, Gln | G |
| A | ATT | Ile | | | ACT | Thr | | | AAT | Asn | | | AGT | Ser | AAT | Asn | T |
| | ATC | Ile | | | ACC | Thr | | | AAC | Asn | | | AGC | Ser | AAC | Asn | C |
| | ATA | Ile | | | ACA | Thr | | | AAA | Lys | | | AGA | Arg | AAA | Lys | A |
| | ATG | Met | ATA | Ile | ACG | Thr | ACA | Thr | AAG | Lys | AAA | Lys | AGG | Arg | AAG, AGA, AAA | Lys, Arg, Lys | G |
| G | GTT | Val | ATT | Ile | GCT | Ala | ACT | Thr | GAT | Asp | AAT | Asn | GGT | Gly | AGT, GAT, AAT | Ser, Asp, Asn | T |
| | GTC | Val | ATC | Ile | GCC | Ala | ACC | Thr | GAC | Asp | AAC | Asn | GGC | Gly | AGC, GAC, AAC | Ser, Asp, Asn | C |
| | GTA | Val | ATA | Ile | GCA | Ala | ACA | Thr | GAA | Glu | AAA | Lys | GGA | Gly | AGA, AAA, GAA | Arg, Lys, Glu | A |
| | GTG | Val | ATG, GTA, ATA | Met, Val, Ile | GCG | Ala | ACG, GCA, ACA | Thr, Ala, Thr | GAG | Glu | AAG, GAA, AAA | Lys, Glu, Lys | GGG | Gly | AGG, GAG, GGA, AAG, GAA, AAG, AAA | Arg, Glu, Gly, Lys, Glu, Lys, Lys | G |
| | From | | To | | From | | To | | From | | To | | From | | To | | |

### Supplementary Table 1c. A-to-G editing

| 1st | 2nd | | | | | | | | | | | | | | | | 3rd |
|---|---|---|---|---|---|---|---|---|---|---|---|---|---|---|---|---|---|
| | T | | | | C | | | | A | | | | G | | | | |
| T | TTT | Phe | | | TCT | Ser | | | TAT | Tyr | TGT | Cys | TGT | Cys | | | T |
| | TTC | Phe | | | TCC | Ser | | | TAC | Tyr | TGC | Cys | TGC | Cys | | | C |
| | TTA | Leu | TTG | Leu | TCA | Ser | TCG | Ser | TAA | STOP | TGA, TAG, TGG | STOP, STOP, Trp | TGA | STOP | TGG | Trp | A |
| | TTG | Leu | | | TCG | Ser | | | TAG | STOP | TGG | Trp | TGG | Trp | | | G |
| C | CTT | Leu | | | CCT | Pro | | | CAT | His | CGT | Arg | CGT | Arg | | | T |
| | CTC | Leu | | | CCC | Pro | | | CAC | His | CGC | Arg | CGC | Arg | | | C |
| | CTA | Leu | CTG | Leu | CCA | Pro | CCG | Pro | CAA | Gln | CGA, CGG, CAG | Arg, Arg, Gln | CGA | Arg | CGG | Arg | A |
| | CTG | Leu | | | CCG | Pro | | | CAG | Gln | CGG | Arg | CGG | Arg | | | G |
| A | ATT | Ile | GTT | Val | ACT | Thr | GCT | Ala | AAT | Asn | GAT, AGT, GGT | Asp, Ser, Gly | AGT | Ser | GGT | Gly | T |
| | ATC | Ile | GTC | Val | ACC | Thr | GCC | Ala | AAC | Asn | GAC, AGC, GGC | Asp, Ser, Gly | AGC | Ser | GGC | Gly | C |
| | ATA | Ile | GTA, ATG, GTG | Val, Met, Val | ACA | Thr | GCA, ACG, GCG | Ala, Thr, Ala | AAA | Lys | GAA, AGA, AAG, GGA, GAG, AGG, GGG | Glu, Arg, Lys, Gly, Glu, Arg, Gly | AGA | Arg | GGA, AGG, GGG | Gly, Arg, Gly | A |
| | ATG | Met | GTG | Val | ACG | Thr | GCG | Ala | AAG | Lys | GAG, AGG, GGG | Glu, Arg, Gly | AGG | Arg | GGG | Gly | G |
| G | GTT | Val | | | GCT | Ala | | | GAT | Asp | GGT | Gly | GGT | Gly | | | T |
| | GTC | Val | | | GCC | Ala | | | GAC | Asp | GGC | Gly | GGC | Gly | | | C |
| | GTA | Val | GTG | Val | GCA | Ala | GCG | Ala | GAA | Glu | GGA, GAG, GGG | Gly, Glu, Gly | GGA | Gly | GGG | Gly | A |
| | GTG | Val | | | GCG | Ala | | | GAG | Glu | GGG | Gly | GGG | Gly | | | G |
| | From | | To | | From | | To | | From | | To | | From | | To | | |

### Supplementary Table 1d. T-to-C editing

| 1st | 2nd | | | | | | | | | | | | | | | | 3rd |
|---|---|---|---|---|---|---|---|---|---|---|---|---|---|---|---|---|---|
| | T | | | | C | | | | A | | | | G | | | | |
| T | TTT | Phe | CTT, TCT, TTC, CCT, CTC, TCC, CCC | Leu, Ser, Phe, Pro, Leu, Ser, Pro | TCT | Ser | CCT, TCC, CCC | Pro, Ser, Pro | TAT | Tyr | CAT, TAC, CAC | His, Tyr, His | TGT | Cys | CGT, TGC, CGC | Arg, Cys, Arg | T |
| | TTC | Phe | CTC, TCC, CCC | Leu, Ser, Pro | TCC | Ser | CCC | Pro | TAC | Tyr | CAC | His | TGC | Cys | CGC | Arg | C |
| | TTA | Leu | CTA, TCA, CCA | Leu, Ser, Pro | TCA | Ser | CCA | Pro | TAA | STOP | CAA | Gln | TGA | STOP | CGA | Arg | A |
| | TTG | Leu | CTG, TCG, CCG | Leu, Ser, Pro | TCG | Ser | CCG | Pro | TAG | STOP | CAG | Gln | TGG | Trp | CGG | Arg | G |
| C | CTT | Leu | CCT, CTC, CCC | Pro, Leu, Pro | CCT | Pro | CCC | Pro | CAT | His | CAC | His | CGT | Arg | CGC | Arg | T |
| | CTC | Leu | CCC | Pro | CCC | Pro | | | CAC | His | | | CGC | Arg | | | C |
| | CTA | Leu | CCA | Pro | CCA | Pro | | | CAA | Gln | | | CGA | Arg | | | A |
| | CTG | Leu | CCG | Pro | CCG | Pro | | | CAG | Gln | | | CGG | Arg | | | G |
| A | ATT | Ile | ACT, ATC, ACC | Thr, Ile, Thr | ACT | Thr | ACC | Thr | AAT | Asn | AAC | Asn | AGT | Ser | AGC | Ser | T |
| | ATC | Ile | ACC | Thr | ACC | Thr | | | AAC | Asn | | | AGC | Ser | | | C |
| | ATA | Ile | ACA | Thr | ACA | Thr | | | AAA | Lys | | | AGA | Arg | | | A |
| | ATG | Met | ACG | Thr | ACG | Thr | | | AAG | Lys | | | AGG | Arg | | | G |
| G | GTT | Val | GCT, GTC, GCC | Ala, Val, Ala | GCT | Ala | GCC | Ala | GAT | Asp | GAC | Asp | GGT | Gly | GGC | Gly | T |
| | GTC | Val | GCC | Ala | GCC | Ala | | | GAC | Asp | | | GGC | Gly | | | C |
| | GTA | Val | GCA | Ala | GCA | Ala | | | GAA | Glu | | | GGA | Gly | | | A |
| | GTG | Val | GCG | Ala | GCG | Ala | | | GAG | Glu | | | GGG | Gly | | | G |
| | From | | To | | From | | To | | From | | To | | From | | To | | |

The original codon table is filled with grey, the generated codons are filled with blue (CBE) and green (ABE). Possible combinations to generate STOP codons by CBE are shown in red.

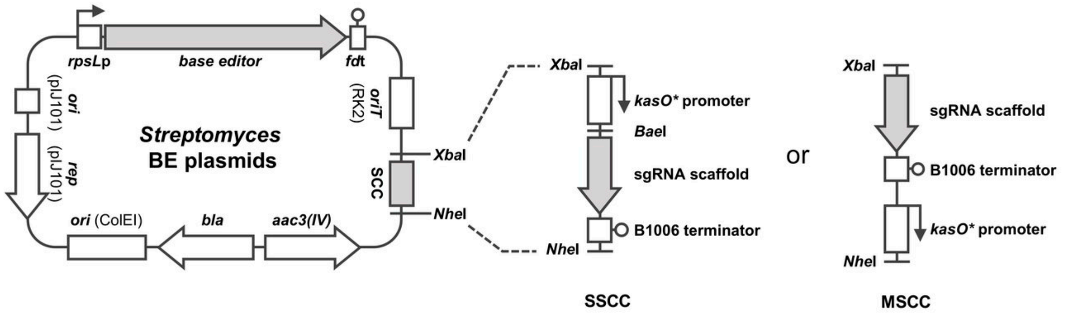


| CBE plasmids | Base editor | Base pair conversion | Efficiency | SCC | Target | Size (bp) |
|---|---|---|---|---|---|---|
| pSCBE2-single | BE2 | C•G → T•A | Medium | SSCC | 1 | 13640 |
| pSCBE3-single | BE3 | C•G → T•A | High | SSCC | 1 | 13640 |
| pSCBE3-HF-single | HF-BE3 | C•G → T•A | High | SSCC | 1 | 13640 |
| pSCBE2 | BE2 | C•G → T•A | Medium | MSCC | 1-n | 13622 |
| pSCBE3 | BE3 | C•G → T•A | High | MSCC | 1-n | 13622 |
| pSCBE3-HF | HF-BE3 | C•G → T•A | High | MSCC | 1-n | 13622 |

| ABE plasmids | Base editor | Base pair conversion | Efficiency | SCC | Target | Size (bp) |
|---|---|---|---|---|---|---|
| pSABEd-single | ABEd | A•T → G•C | Medium | SSCC | 1 | 13835 |
| pSABEn-single | ABEn | A•T → G•C | High | SSCC | 1 | 13835 |
| pSABEd | ABEd | A•T → G•C | Medium | MSCC | 1-n | 13817 |
| pSABEn | ABEn | A•T → G•C | High | MSCC | 1-n | 13817 |

**Supplementary Fig. 2** Schematic of the *Streptomyces* base editing plasmids. *ori(ColE1)*, *E. coli* replicon; *ori(pIJ101)* and *rep(pIJ101)*, *Streptomyces* replicon; *oriT(RK2)*, origin of transfer derived from RK2 plasmid; *bla*, ampicillin resistance gene; *aac(3)IV*, apramycin resistance gene.

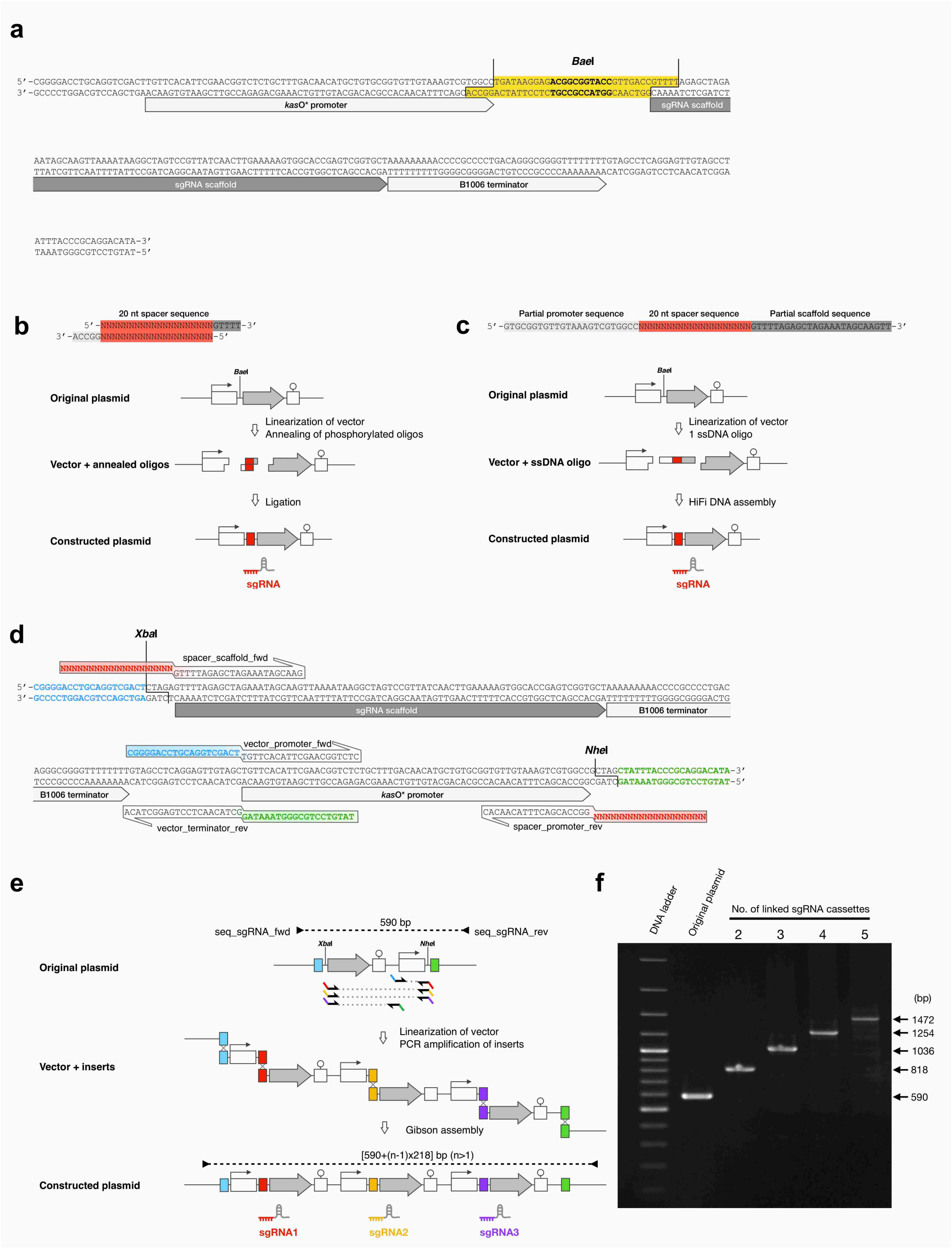


**Supplementary Fig. 3** Schematics of the sgRNA cloning strategy, related to Fig. 1c. **a** Complete sequence of SSCC. The type-IIS restriction endonuclease site *Bae*I removes the yellow highlighted region from the vector. **b** Cloning of the 20-nt spacer into SSCC by annealing of phosphorylated DNA oligos. **c** Cloning of the 20-nt spacer into SSCC by Hifi DNA assembly of single strand DNA oligo. **d** Complete sequence of MSCC. The MSCC serves as PCR template for the amplification of Gibson assembly inserts. The 20-nt spacer sequences could be introduced by primers. **e** Rearrangement of the elements is required to construct the functional sgRNA expression cassettes (*kasO*p*/20-nt spacer/sgRNA scaffold/B1006t). MSCC enables the cloning of single or multiple sgRNA, the schematic diagram shows the coning of 3 sgRNA. **f** PCR confirmation of the recombinants harboring 2 to 5 sgRNA cloned by MSCC. Vent DNA polymerase (NEB); primer pair: seq_sgRNA_fwd and seq_sgRNA_rev; 1.7% agarose; GeneRuler 100 bp Plus DNA Ladder (ThermoFisher).

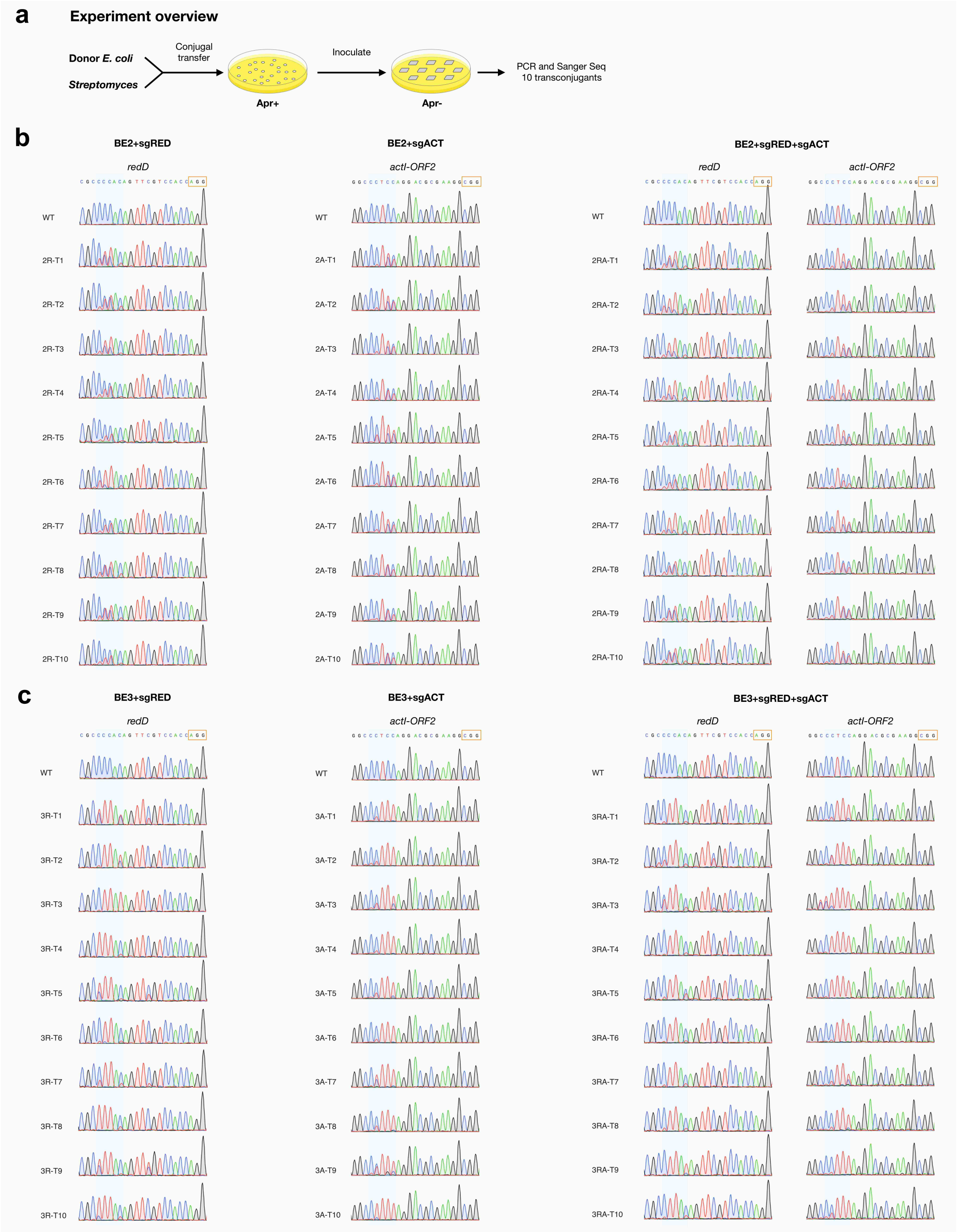


**Supplementary Fig. 4** Verification of CBE mediated C-to-T editing in single transconjugants of *S. coelicolor*, related to Fig. 2b. **a** Experiment overview after conjugal transfer. DNA sample is from the individual transconjugants. **b** Sanger sequencing result of BE2-derived transconjugants. **c** Sanger sequencing result of BE3-derived transconjugants.

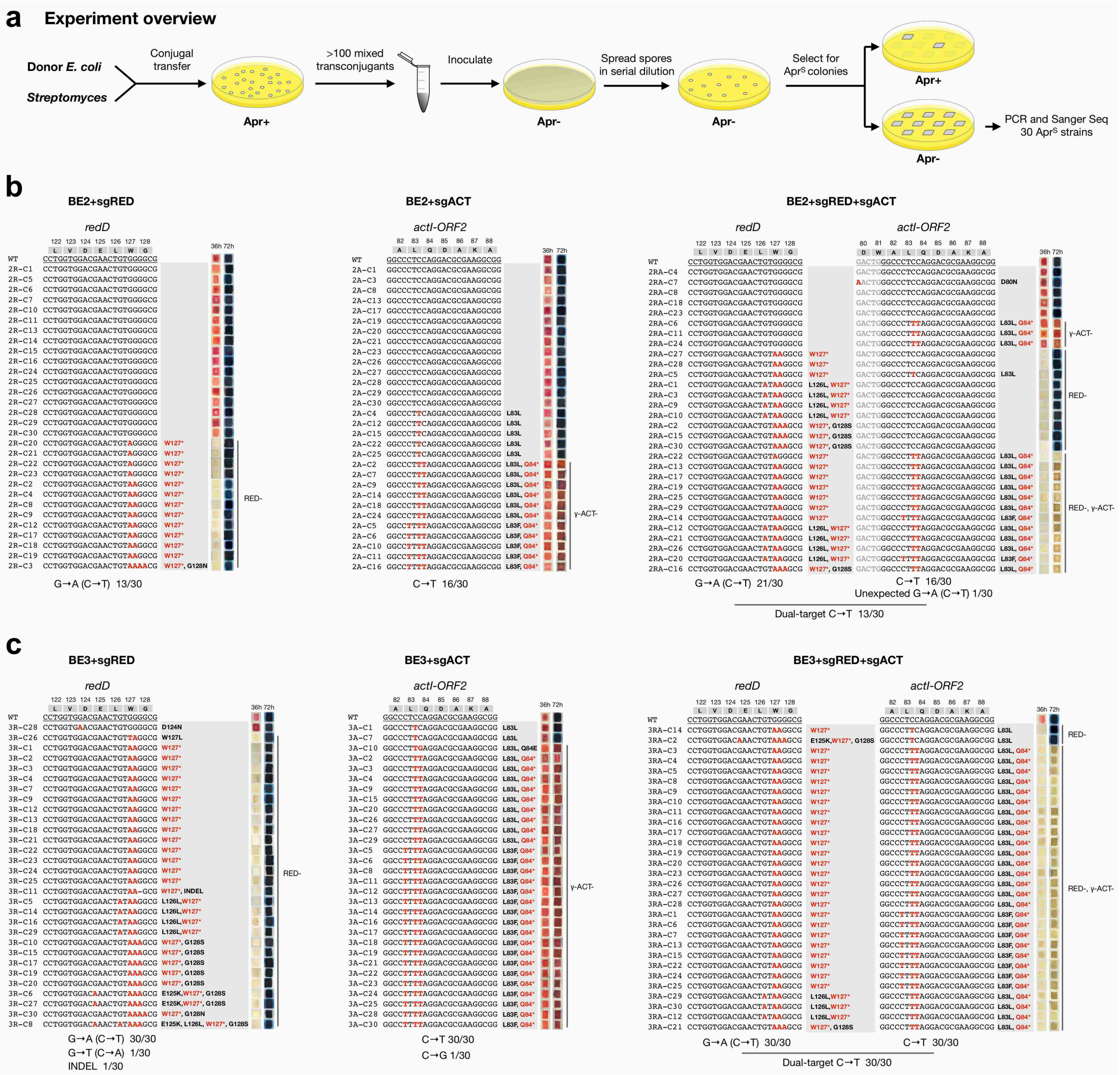


**Supplementary Fig. 5** Geno- and phenotyping of the *S. coelicolor* mutants after CBE editing and plasmid curing, related to Fig. 2c. **a** Experiment overview after conjugal transfer. DNA sample is from the plasmid-free colonies. **b** Regenerated mutants after BE2 editing. **c** Regenerated mutants after BE3 editing.

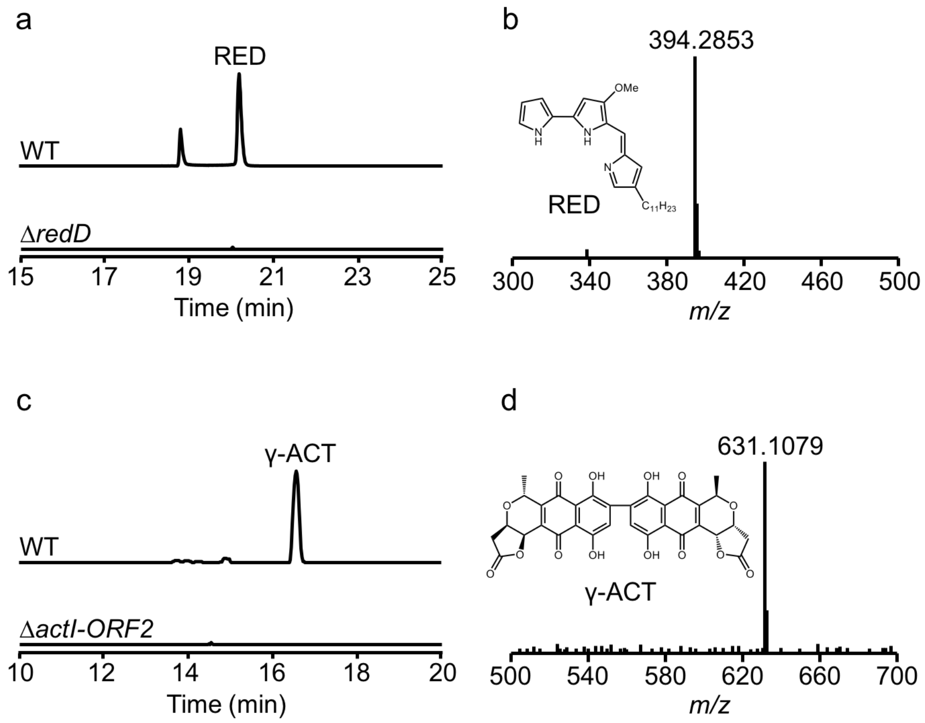


**Supplementary Fig. 6** LC-ESI-HRMS analysis of RED (a-b) and γ-ACT (c-d) in wild type *S. coelicolor* and its mutants, related to Fig. 2c.

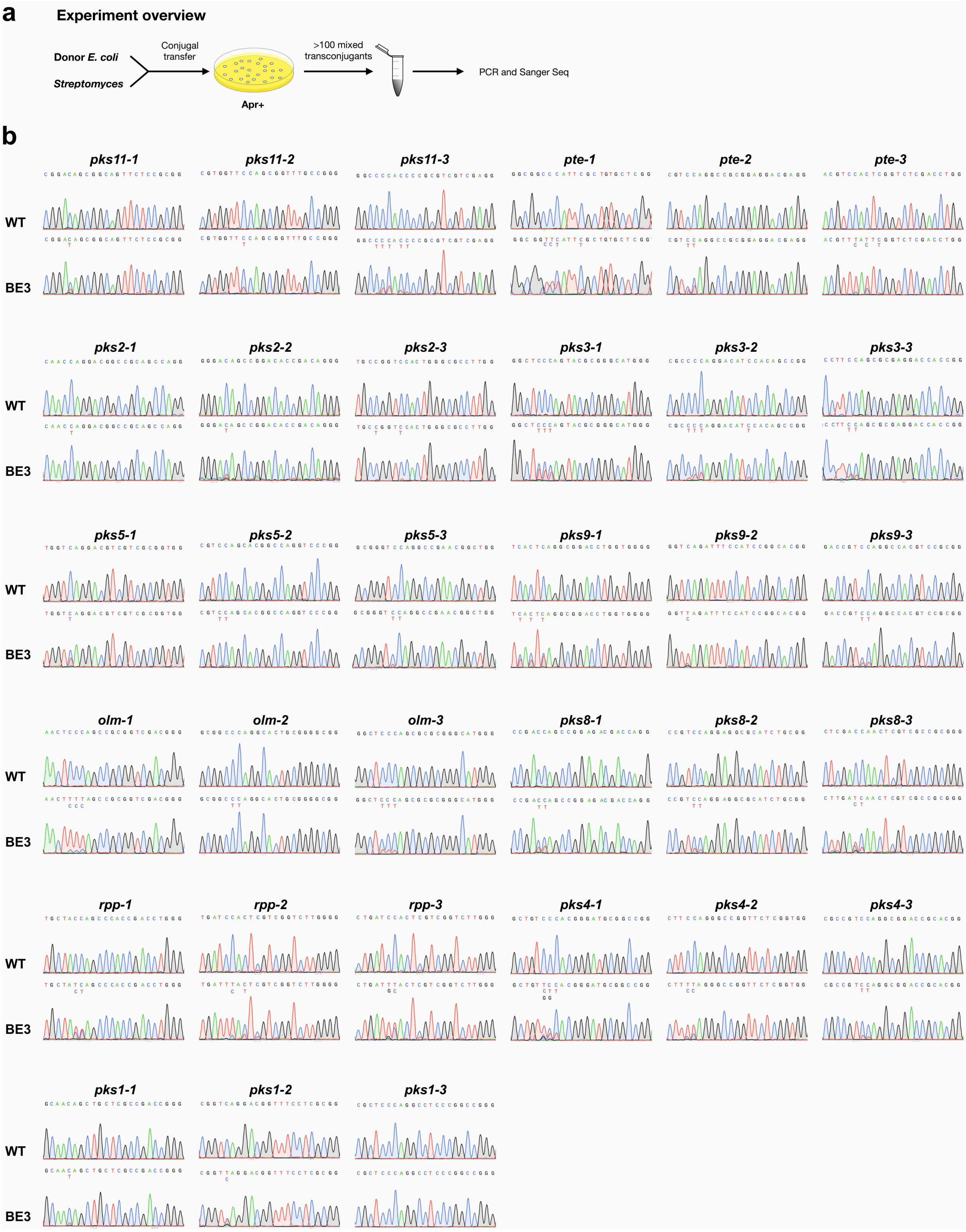


**Supplementary Fig. 7** BE3 mediated C-to-T editing in *S. avermitilis*. **a** Experiment overview after conjugal transfer. The 33 targets are examined individually. For each target, DNA sample is from the mixture of over 100 transconjugants. **b** Sanger sequencing result of the 33 targets.

**Supplementary Table 2** Summary of the target sequence and editing efficiency collected from Supplementary Fig. 7.

| sgRNA name | Spacer | PAM | Spacer G+C content (%) | C-to-T conversion (%) | | | | | Applied to multiplexing |
|---|---|---|---|---|---|---|---|---|---|
| | | | | C4 | C5 | C6 | C7 | C8 | |
| sgPKS11-1 | CGGACAGCGGCAGTTCTCCG | CGG | 70 | - | 28.7 | - | - | 2.0 | ✔ |
| sgPKS11-2 | CGTGGTTCCAGCGGTTTGCC | GGG | 65 | - | - | - | - | 21.8 | |
| sgPKS11-3 | GGCCCCACCCCGCGTCGTCG | AGG | 85 | 7.1 | 29.5 | 23.5 | - | 16.7 | |
| sgPTE-1 | GGCGGCCCATTCGCTGTGCT | CGG | 70 | - | - | 61.8 | 58.9 | 35.8 | |
| sgPTE-2 | CGTCCAGGCCGCGGAGGACG | AGG | 80 | 36.4 | 34.1 | - | - | - | |
| sgPTE-3 | ACGTCCACTCGGTCTCGACC | TGG | 65 | - | 95.4 | 84.8 | - | 71.0 | ✔ |
| sgPKS2-1 | CAACCAGGACGGCCGCAGCC | AGG | 75 | 3.3 | 3.3 | - | - | | |
| sgPKS2-2 | GGGACAGCCGGACACCGACA | GGG | 70 | - | 21.1 | - | - | 0.0 | ✔ |
| sgPKS2-3 | TGCCGGTCCACTGGGCGCCT | TGG | 75 | 7.6 | - | - | - | 9.9 | |
| sgPKS3-1 | GGCTCCCAGTACGCGGGCAT | GGG | 70 | - | 38.9 | 35.1 | 35.9 | - | ✔ |
| sgPKS3-2 | CGCCCCAGGACATCCACAGC | CGG | 70 | 12.7 | 29.7 | 23.5 | - | - | |
| sgPKS3-3 | CCTTCCAGCGCGAGGACCAC | CGG | 70 | - | 28.3 | 21.9 | - | - | |
| sgPKS5-1 | TGGTCAGGACGTCGTCGCGG | TGG | 70 | - | 33.3 | - | - | - | ✔ |
| sgPKS5-2 | CGTCCAGCACGGCCAGGTCC | CGG | 75 | 7.5 | 6.7 | - | - | 0.0 | |
| sgPKS5-3 | GCGGGTCCAGGCCGAACGGC | TGG | 80 | - | - | - | 19.2 | 4.5 | |
| sgPKS9-1 | TCACTCAGGCGGACCTGGTG | GGG | 65 | 32.7 | - | 43.9 | - | - | |
| sgPKS9-2 | GGTCAGATTTCCATCCGGCA | CGG | 55 | 69.7 | - | - | - | - | ✔ |
| sgPKS9-3 | GACCGTCCAGGCCACGTCCG | CGG | 75 | 2.8 | - | - | 29.6 | 19.7 | |
| sgOLM-1 | AACTCCCAGCCGCGGTCGAC | GGG | 70 | - | 84.6 | 85.9 | 79.0 | - | ✔ |
| sgOLM-2 | GCGGCCCAGGCACTGCGGGG | CGG | 85 | - | 1.6 | 4.0 | 2.6 | - | |
| sgOLM-3 | GGCTCCCAGCGCGCGGGCAT | GGG | 80 | - | 23.8 | 17.3 | 15.6 | - | |
| sgPKS8-1 | CCGACCAGCCGGAGACGACC | AGG | 75 | - | 30.6 | 30.4 | - | - | ✔ |
| sgPKS8-2 | CCGTCCAGGAGGCGCATCTG | CGG | 70 | - | 23.3 | 18.4 | - | - | |
| sgPKS8-3 | CTCGACCAACTCGTCGCCGC | GGG | 70 | - | - | 51.1 | 22.5 | - | |
| sgRPP-1 | TGCTACCAGCCCACCGACCT | GGG | 65 | - | - | 63.9 | 46.5 | - | |
| sgRPP-2 | TGATCCACTCGTCGGTCTTG | GGG | 55 | - | 88.9 | 81.4 | - | 25.9 | ✔ |
| sgRPP-3 | CTGATCCACTCGTCGGTCTT | GGG | 55 | - | - | 73.7 | 57.1 | - | |
| sgPKS4-1 | GCTGTCCCACGGGATGCGGC | CGG | 75 | - | - | 60.8 | 33.8 | 19.5 | |
| sgPKS4-2 | CTTCCAGGGCCGGTTCTCGG | TGG | 70 | 74.1 | 67.0 | - | - | - | ✔ |
| sgPKS4-3 | CGCCGTCCAGGCGGACCGCA | CGG | 80 | 2.1 | | - | 19.9 | 15.3 | |
| sgPKS1-1 | GCAACAGCTGCTCGCCGACC | GGG | 70 | - | 14.5 | - | - | 2.1 | |
| sgPKS1-2 | CGGTCAGGACGGTTTCCTCG | CGG | 65 | - | 41.2 | - | - | - | ✔ |
| sgPKS1-3 | CGCTCCCAGGCCTCCCGGCC | GGG | 85 | - | 2.2 | 0.7 | 0.9 | - | |

The 4-8 nt editing widow is shadowed in blue. Cs located in editing window are bolded. Codons to generate premature stop codons are underlined.

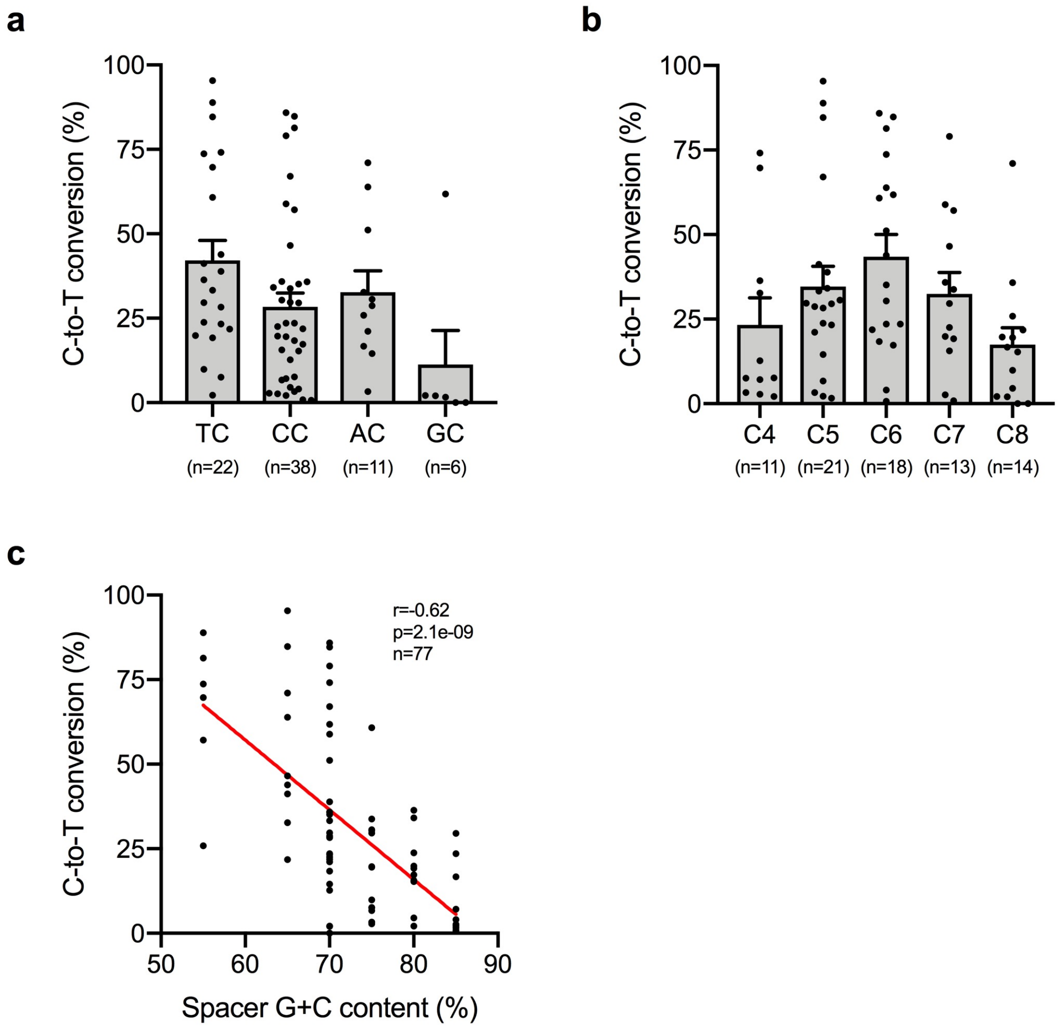


**Supplementary Fig. 8** Analysis of sequence context and editing efficiency using data from Supplementary Table 2. **a**-**b** Effect from the 5'-motif and position. Values and error bars represent mean ± s.e.m. **c** Correlation between editing efficiency and spacer G+C content.

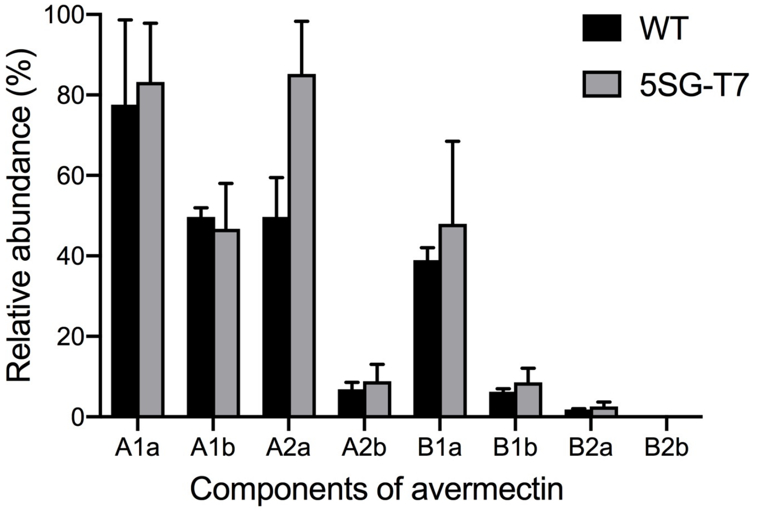


**Supplementary Fig. 9** LC-ESI-HRMS analysis of eight avermectin components production in wild type *S. avermitilis* and the mutant 5SG-T7. Peak area was calculated to represent production. Values and error bars represent mean ± s.d. of 3 repeats.

**Supplementary Table 3** Potential off-target sites (OTs) in the *ave* gene cluster associated with the sgRNAs targeting the 11 PKS gene clusters.

| Name | Location | Site | No. of mismatches |
|---|---|---|---|
| OLM-ON | SAV_Chr:3623187-3623210:+ | AACTCCCA_GCCGCGGTCGAC-GGG | 0 |
| OLM-OT-1 | SAV_Chr:1144179-1144202:- | AggTCCCA_GCCGCGGTCGgt-GGG | 4 |
| OLM-OT-2 | SAV_Chr:1157931-1157954:- | AggTCCCA_GCCGCGGTCGgt-GGG | 4 |
| OLM-OT-3 | SAV_Chr:1162704-1162727:- | AggTCCCA_GCCGCGGTCGgt-CGG | 4 |
| OLM-OT-4 | SAV_Chr:1194538-1194561:+ | AcgTCCCA_aCCcCGGTCGAC-CGG | 4 |
| OLM-OT-5 | SAV_Chr:1197673-1197696:+ | AcgTCCCA_aCCcCGGTCGAC-CGG | 4 |
| OLM-OT-6 | SAV_Chr:1202344-1202367:+ | AcgTCCCA_aCCcCGGTCGAC-CGG | 4 |
| PKS4-ON | SAV_Chr:8556845-8556868:+ | CTTCCAGG_GCCGGTTCTCGG-TGG | 0 |
| PKS4-OT | SAV_Chr:1166353-1166376:- | CgcCCAGG_GCCGGTcCgCGG-TGG | 4 |
| PKS1-ON | SAV_Chr:8789027-8789050:- | CGGTCAGG_ACGGTTTCCTCG-CGG | 0 |
| PKS1-OT | SAV_Chr:1132331-1132354:- | CGGTgcGG_gCGGTgTCCTCG-GGG | 4 |

The OTs were predicted by CasOT, with the selection criteria of up to two mismatches in seed region and up to four mismatches in total. Sequences are showed in the form of <non-seed>_<seed>-<PAM>. Mismatched bases are showed in lower case.

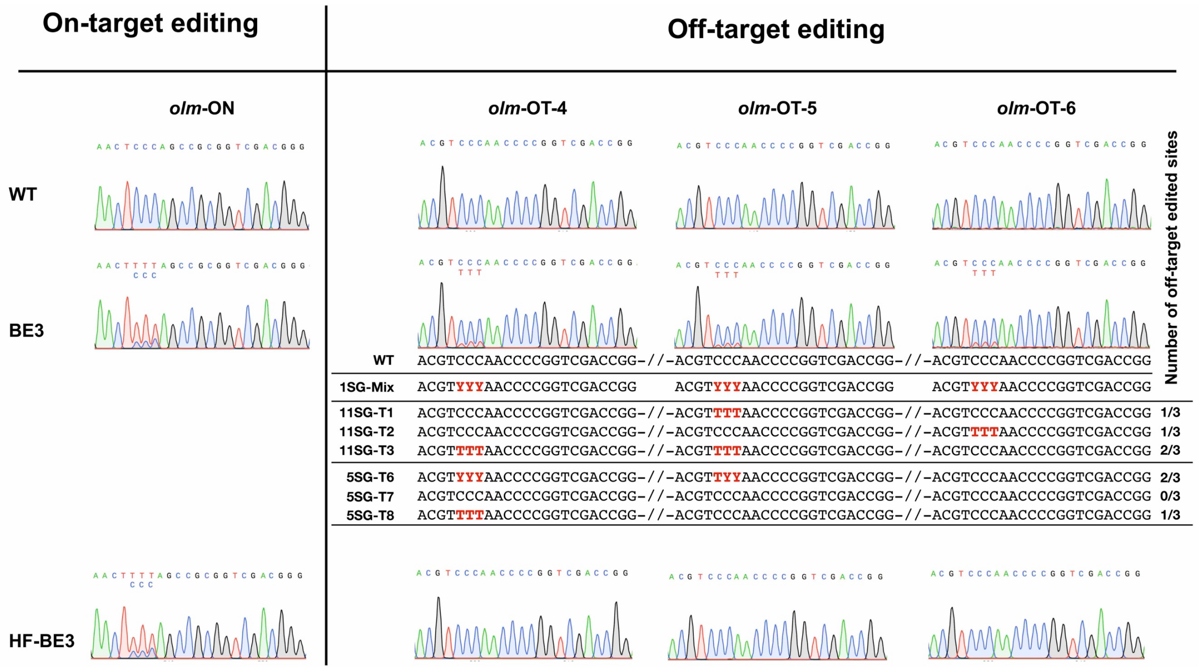


**Supplementary Fig. 10** On- and off-target editing of BE3 and HF-BE3 in *S. avermitilis* using sgOLM-1, related to Fig. 3 and Fig. 4. Sanger sequencing chromatograms represent WT and the editing event in the mixture of over 100 transconjugants that accepting the only sgOLM-1 containing BE plasmids. Alignment show the off-target editing in the mutants that accepting the 5- or 11-sgRNA array combined BE3 plasmids.

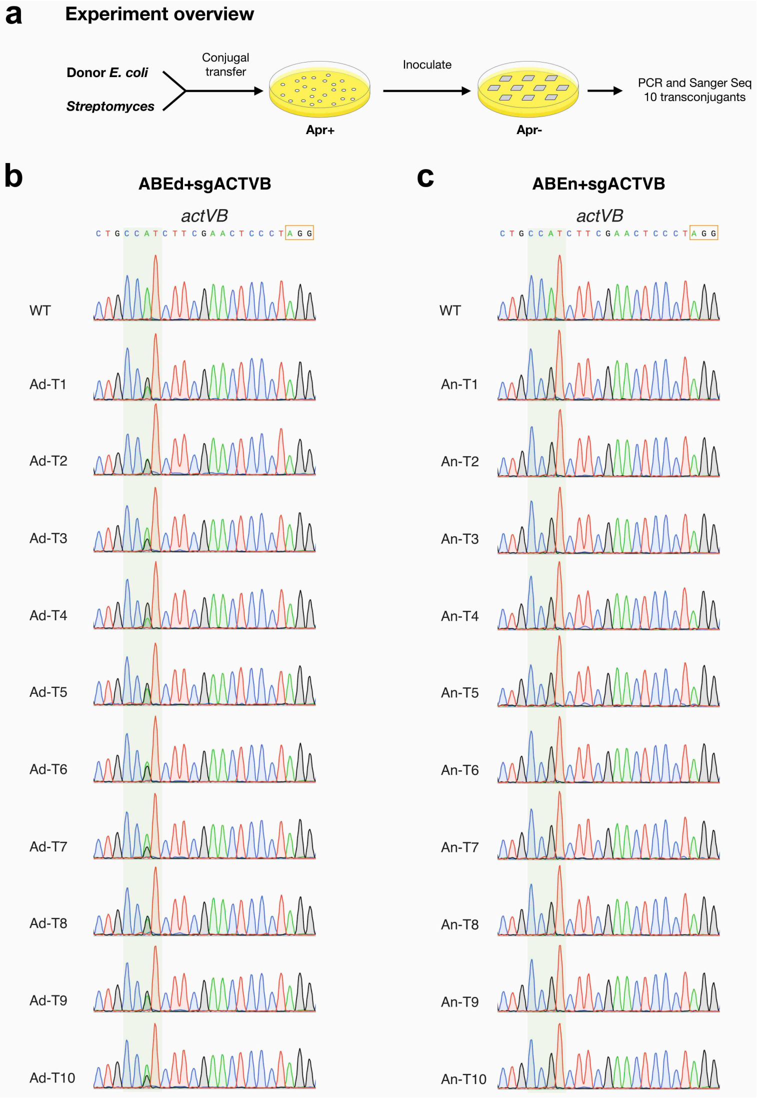


**Supplementary Fig. 11** Verification of ABE mediated A-to-G editing in single transconjugants of *S. coelicolor*, related to Fig. 5b. **a** Experiment overview after conjugal transfer. DNA sample is from the individual transconjugants. **b** Sanger sequencing result of ABEd-derived transconjugants. **c** Sanger sequencing result of ABEn-derived transconjugants.

**Supplementary Table 4** Primers used in this study.

| Name | Sequence (5'-3') |
| --- | --- |
| pYH7-REMOVE-COS-fwd | acacgtctgaagctAGCTATTTACCCGCAGGACATATCCA |
| pYH7-REMOVE-COS-rev | GGCGAAAAGCCGAGCTCATCGGTCAGCTTCTCAACC |
| INSERT-Cas9-fwd | CAGCATCGGCCTGGCCATCGGCACCAACAGCGT |
| CBE-INSERT-Cas9-rev | tggtggagccgcccgaGTCGCCGCCCAGCTGGCTC |
| ABE-INSERT-Cas9-rev | cgtagatctgaattctcaGTCGCCGCCCAGCTGGCTC |
| Cas9-H840A-fwd | GACTACGACGTCGACGCCATCGTGCCGCAGTCC |
| Cas9-H840A-rev | ACTGCGGCACGATGGCGTCGACGTCGTAGTCGG |
| ICE-sgRNA-anti | AAAAGCACCGACTCGGTGCCACTTTTTCAAGTTGATAACG<br>GACTAGCCTTATTTTAACTTGCTATTTCTAGCTCTAAAAC |
| ICE-pSCBE3-1 | GATCACTAATACGACTCACTATAGCTCGTCAAGCTGAACCGCG<br>GTTTTAGAGCTAGAAA |
| ICE-pSCBE3-2 | GATCACTAATACGACTCACTATACCACGTGCTTGGTGATCTGC<br>GTTTTAGAGCTAGAAA |
| HF-Cas9-fwd | GGAGCTGCTCGTCAAGCTGAACC |
| HF-Cas9-N497A-rev | CTTGTCGAAGGCGGTCATGCGCTCGATGAACGAC |
| HF-Cas9-N497A-fwd | GCGCATGACCGCCTTCGACAAGAACCTCCCGAACG |
| HF-Cas9-R661A-rev | CCGGGAGAGGGCGCCCCAGCCGGTGTACCG |
| HF-Cas9-R661A-fwd | CGGCTGGGGCGCCCTCTCCCGGAAGCTGATCAACG |
| HF-Cas9-Q695A-rev | GTCGTGGATGAGCGCCATGAAGTTGCGGTTGGCG |
| HF-Cas9-Q695A-fwd | CAACTTCATGGCGCTCATCCACGACGACAGCCTG |
| HF-Cas9-Q929A-rev | GGCCACGTGCTTGGTGATCGCCCGGGTCTCGACCAG |
| vector-promoter-fwd | cggggacctgcaggtcgactTGTTCACATTCGAACGGTCTC |
| vector-terminator-rev | tatgtcctgcgggtaaatagGCTACAACTCCTGAGGCTACA |
| BaeI_spacer-scaffold-fwd | tgataaggagacggcggtaccgttgaccGTTTTAGAGCTAGAAATAGCAAG |
| BaeI_spacer-promoter-rev | ggtcaacggtaccgccgtctccttatcaGGCCACGACTTTACAAC |
| actI2_spacer-scaffold-fwd | ggccctccaggacgcgaaggGTTTTAGAGCTAGAAATAGCAAG |
| actI2_spacer-promoter-rev | ccttcgcgtcctggagggccGGCCACGACTTTACAAC |
| redD_spacer-scaffold-fwd | cgccccacagttcgtccaccGTTTTAGAGCTAGAAATAGCAAG |
| redD_spacer-promoter-rev | ggtggacgaactgtggggcgGGCCACGACTTTACAAC |
| pks11-1_spacer-scaffold-fwd | cggacagcggcagttctccgGTTTTAGAGCTAGAAATAGCAAG |
| pks11-1_spacer-promoter-rev | cggagaactgccgctgtccgGGCCACGACTTTACAAC |
| pks11-2_spacer-scaffold-fwd | cgtggttccagcggtttgccGTTTTAGAGCTAGAAATAGCAAG |
| pks11-2_spacer-promoter-rev | ggcaaaccgctggaaccacgGGCCACGACTTTACAAC |
| pks11-3_spacer-scaffold-fwd | ggccccaccccgcgtcgtcgGTTTTAGAGCTAGAAATAGCAAG |
| pks11-3_spacer-promoter-rev | cgacgacgcggggtggggccGGCCACGACTTTACAAC |
| pte-1_spacer-scaffold-fwd | ggcggcccattcgctgtgctGTTTTAGAGCTAGAAATAGCAAG |
| pte-1_spacer-promoter-rev | agcacagcgaatgggccgccGGCCACGACTTTACAAC |

| | |
|---|---|
| pte-2_spacer-scaffold-fwd | cgtccaggccgcggaggacgGTTTTAGAGCTAGAAATAGCAAG |
| pte-2_spacer-promoter-rev | cgtcctccgcggcctggacgGGCCACGACTTTACAAC |
| pte-3_spacer-scaffold-fwd | acgtccactcggtctcgaccGTTTTAGAGCTAGAAATAGCAAG |
| pte-3_spacer-promoter-rev | ggtcgagaccgagtggacgtGGCCACGACTTTACAAC |
| pks2-1_spacer-scaffold-fwd | caaccaggacggccgcagccGTTTTAGAGCTAGAAATAGCAAG |
| pks2-1_spacer-promoter-rev | ggctgcggccgtcctggttgGGCCACGACTTTACAAC |
| pks2-2_spacer-scaffold-fwd | gggacagccggacaccgacaGTTTTAGAGCTAGAAATAGCAAG |
| pks2-2_spacer-promoter-rev | tgtcggtgtccggctgtcccGGCCACGACTTTACAAC |
| pks2-3_spacer-scaffold-fwd | tgccggtccactgggcgcctGTTTTAGAGCTAGAAATAGCAAG |
| pks2-3_spacer-promoter-rev | aggcgcccagtggaccggcaGGCCACGACTTTACAAC |
| pks3-1_spacer-scaffold-fwd | ggctcccagtacgcgggcatGTTTTAGAGCTAGAAATAGCAAG |
| pks3-1_spacer-promoter-rev | atgcccgcgtactgggagccGGCCACGACTTTACAAC |
| pks3-2_spacer-scaffold-fwd | cgccccaggacatccacagcGTTTTAGAGCTAGAAATAGCAAG |
| pks3-2_spacer-promoter-rev | gctgtggatgtcctggggcgGGCCACGACTTTACAAC |
| pks3-3_spacer-scaffold-fwd | ccttccagcgcgaggaccacGTTTTAGAGCTAGAAATAGCAAG |
| pks3-3_spacer-promoter-rev | gtggtcctcgcgctggaaggGGCCACGACTTTACAAC |
| pks5-1_spacer-scaffold-fwd | tggtcaggacgtcgtcgcggGTTTTAGAGCTAGAAATAGCAAG |
| pks5-1_spacer-promoter-rev | ccgcgacgacgtcctgaccaGGCCACGACTTTACAAC |
| pks5-2_spacer-scaffold-fwd | cgtccagcacggccaggtccGTTTTAGAGCTAGAAATAGCAAG |
| pks5-2_spacer-promoter-rev | ggacctggccgtgctggacgGGCCACGACTTTACAAC |
| pks5-3_spacer-scaffold-fwd | gcgggtccaggccgaacggcGTTTTAGAGCTAGAAATAGCAAG |
| pks5-3_spacer-promoter-rev | gccgttcggcctggacccgcGGCCACGACTTTACAAC |
| pks9-1_spacer-scaffold-fwd | tcactcaggcggacctggtgGTTTTAGAGCTAGAAATAGCAAG |
| pks9-1_spacer-promoter-rev | caccaggtccgcctgagtgaGGCCACGACTTTACAAC |
| pks9-2_spacer-scaffold-fwd | ggtcagatttccatccggcaGTTTTAGAGCTAGAAATAGCAAG |
| pks9-2_spacer-promoter-rev | tgccggatggaaatctgaccGGCCACGACTTTACAAC |
| pks9-3_spacer-scaffold-fwd | gaccgtccaggccacgtccgGTTTTAGAGCTAGAAATAGCAAG |
| pks9-3_spacer-promoter-rev | cggacgtggcctggacggtcGGCCACGACTTTACAAC |
| olm-1_spacer-scaffold-fwd | aactcccagccgcggtcgacGTTTTAGAGCTAGAAATAGCAAG |
| olm-1_spacer-promoter-rev | gtcgaccgcggctgggagttGGCCACGACTTTACAAC |
| olm-2_spacer-scaffold-fwd | gcggcccaggcactgcggggGTTTTAGAGCTAGAAATAGCAAG |
| olm-2_spacer-promoter-rev | ccccgcagtgcctgggccgcGGCCACGACTTTACAAC |
| olm-3_spacer-scaffold-fwd | ggctcccagcgcgcgggcatGTTTTAGAGCTAGAAATAGCAAG |
| olm-3_spacer-promoter-rev | atgcccgcgcgctgggagccGGCCACGACTTTACAAC |
| pks8-1_spacer-scaffold-fwd | ccgaccagccggagacgaccGTTTTAGAGCTAGAAATAGCAAG |
| pks8-1_spacer-promoter-rev | ggtcgtctccggctggtcggGGCCACGACTTTACAAC |
| pks8-2_spacer-scaffold-fwd | ccgtccaggaggcgcatctgGTTTTAGAGCTAGAAATAGCAAG |
| pks8-2_spacer-promoter-rev | cagatgcgcctcctggacggGGCCACGACTTTACAAC |
| pks8-3_spacer-scaffold-fwd | ctcgaccaactcgtcgccgcGTTTTAGAGCTAGAAATAGCAAG |
| pks8-3_spacer-promoter-rev | gcggcgacgagttggtcgagGGCCACGACTTTACAAC |

| | |
|---|---|
| rpp-1_spacer-scaffold-fwd | tgctaccagcccaccgacctGTTTTAGAGCTAGAAATAGCAAG |
| rpp-1_spacer-promoter-rev | aggtcggtgggctggtagcaGGCCACGACTTTACAAC |
| rpp-2_spacer-scaffold-fwd | tgatccactcgtcggtcttgGTTTTAGAGCTAGAAATAGCAAG |
| rpp-2_spacer-promoter-rev | caagaccgacgagtggatcaGGCCACGACTTTACAAC |
| rpp-3_spacer-scaffold-fwd | ctgatccactcgtcggtcttGTTTTAGAGCTAGAAATAGCAAG |
| rpp-3_spacer-promoter-rev | aagaccgacgagtggatcagGGCCACGACTTTACAAC |
| pks4-1_spacer-scaffold-fwd | gctgtcccacgggatgcggcGTTTTAGAGCTAGAAATAGCAAG |
| pks4-1_spacer-promoter-rev | gccgcatcccgtgggacagcGGCCACGACTTTACAAC |
| pks4-2_spacer-scaffold-fwd | cttccagggccggttctcggGTTTTAGAGCTAGAAATAGCAAG |
| pks4-2_spacer-promoter-rev | ccgagaaccggccctggaagGGCCACGACTTTACAAC |
| pks4-3_spacer-scaffold-fwd | cgccgtccaggcggaccgcaGTTTTAGAGCTAGAAATAGCAAG |
| pks4-3_spacer-promoter-rev | tgcggtccgcctggacggcgGGCCACGACTTTACAAC |
| pks1-1_spacer-scaffold-fwd | gcaacagctgctcgccgaccGTTTTAGAGCTAGAAATAGCAAG |
| pks1-1_spacer-promoter-rev | ggtcggcgagcagctgttgcGGCCACGACTTTACAAC |
| pks1-2_spacer-scaffold-fwd | cggtcaggacggtttcctcgGTTTTAGAGCTAGAAATAGCAAG |
| pks1-2_spacer-promoter-rev | cgaggaaaccgtcctgaccgGGCCACGACTTTACAAC |
| pks1-3_spacer-scaffold-fwd | cgctcccaggcctcccggccGTTTTAGAGCTAGAAATAGCAAG |
| pks1-3_spacer-promoter-rev | ggccgggaggcctgggagcgGGCCACGACTTTACAAC |
| actVB_spacer-scaffold-fwd | ctgccatcttcgaactccctGTTTTAGAGCTAGAAATAGCAAG |
| actVB_spacer-promoter-rev | agggagttcgaagatggcagGGCCACGACTTTACAAC |
| seq-sgRNA-fwd | CATGCGCTCCATCAAGAAGAGC |
| seq-sgRNA-rev | CGCTGATGATATGCTGACGCTC |
| seq-redD-fwd | CTTCTTCTCTGCCCTCTGACCG |
| seq-redD-rev | GGTCGATCTCCAGCAGGTAGC |
| seq-actI2-fwd | GAAGCTGCGGTCCGTACTCA |
| seq-actI2-rev | TCGAAGGTGGAGGCGCAG |
| seq-pte-1-rev | CCCGGTGTTCGAAGGCGGTG |
| seq-pte-2-fwd | GCGAGCTCGCCTTCCTCTTC |
| seq-pte-2-rev | GGAATGGAAGGCGTGGCTG |
| seq-pte-3-fwd | CGGCATCGTCGTCCTCAAACCTC |
| seq-pte-3-rev | TCCCAGCCACGGTCCGTCGG |
| seq-pks2-1-fwd | GACCGCCTGGGAGACGATC |
| seq-pks2-1-rev | TACCGTGCGCCTCGACGTA |
| seq-pks2-2_3-fwd | CCGGAAAGACCCTCTTCCCG |
| seq-pks2-2_3-rev | TCATCACGGCGAAGAGGACC |
| seq-pks3-1_2_3-fwd | GCTGTGCTACACCGCCAAC |
| seq-pks3-1_2_3-rev | CCGTGAACTCCCGTACCCC |
| seq-pks5-1_2-fwd | CCACCCTTCCCGAGAGCAG |
| seq-pks5-1-rev | GAGGACAGGGTGAAGCTGTC |
| seq-pks5-2-rev | CGCGAGCACAAGATGGCAG |

| | |
|---|---|
| seq-pks5-3-fwd | GTCCCCTGCCGAGGAGAAC |
| seq-pks5-3-rev | GTCGGACCACACGATCTCC |
| seq-pks9-1-fwd | GCGGCGATGTAGAGTCCGT |
| seq-pks9-1-rev | TCACCTCCACCATCGCGAC |
| seq-pks9-2-fwd | GACCAGGTCGGCATCGTCA |
| seq-pks9-2-rev | GGTCGAAGGTGGTGGCGTA |
| seq-pks9-3-fwd | AGTGCGATTTCGACCCGGT |
| seq-pks9-3-rev | GCCTTGATCGCGTCGAAGC |
| seq-olm-1_2-fwd | ATCGTCGGTATGGGCTGCC |
| seq-olm-1_2-rev | CCAGCATCCCGACACCCTC |
| seq-olm-3-fwd | CCGGATTCCTGGCCCACTT |
| seq-olm-3-rev | CGTCAACTGCTTGGTCCGC |
| seq-pks8-1-fwd | GCCCTGGAGTCCCATGTCC |
| seq-pks8-1-rev | CCGTTCGTAGGGTCCGAGG |
| seq-pks8-2-fwd | CTCGGTGGAGCTGTCCTGG |
| seq-pks8-2-rev | ATGTACGGGGAGGGGTCGA |
| seq-pks8-3-fwd | TGGTCACCAGCATCGGCAT |
| seq-pks8-3-rev | GTGAACGTCTTGCGGCAGG |
| seq-rpp-1_2_3-fwd | TCGTGCACGGGGTTCATGA |
| seq-rpp-1_2_3-rev | GGCACTCGCAGGAACTTGC |
| seq-pks4-1_2_3-fwd | CGTCGCACGACAAGGAACG |
| seq-pks4-1_2_3-rev | TTGAGCAGTGCGTCCAGGT |
| seq-pks1-1_2-fwd | CGACAACGATGCCTTTCCCG |
| seq-pks1-1_2-rev | CACGTGCAGGAAGGTGGAGA |
| seq-pks1-3-fwd | GCGCAAGTTCGTCTTCGAGG |
| seq-pks1-3-rev | GCCGTTCAGGTCGTGGTAGA |
| seq-olm-OT-1-fwd | CTGTCTCGCTGGTCTTCGAC |
| seq-olm-OT-1-rev | CGTGTACGAGATACGACCGG |
| seq-olm-OT-2-fwd | TCTGCTCTCCGAGATCGCG |
| seq-olm-OT-2-rev | GATGTAGGAGATTCGGCCGG |
| seq-olm-OT-3-fwd | CTAGACGACGAAGAGGACGC |
| seq-olm-OT-3-rev | CCCTCGATGGTCTCCACATC |
| seq-olm-OT-4-fwd | GAGACGACGCTGTTGGAGAC |
| seq-olm-OT-4-rev | CTCGGTTTCGACTCCCTCAC |
| seq-olm-OT-5-fwd | GGACACCTCCAGCATCAACC |
| seq-olm-OT-5-rev | CTTCTCGACCTGGTCCGTAC |
| seq-olm-OT-6-fwd | GCGTAGTCCTGGGACATGAG |
| seq-olm-OT-6-rev | TCTCAAGCGCGTTACTGCCG |
| seq-pks4-OT-fwd | GTTCCGTGTTGCGCGTACGG |
| seq-pks4-OT-rev | GCAGCCGGCTGTCTGCGAG |

| | |
|---|---|
| seq-pks1-OT-fwd | GGAGTTTCCTGTCTGCACC |
| seq-pks1-OT-rev | GTGACGGTCAGGCAGTAGTC |
| seq-actVB-fwd | GACGCCAACCGGGACTATCTG |
| seq-actVB-rev | CTCCATCGAGACGGACACGAAC |

The mutations to generate Cas9 variants are underlined. The 5'-overhangs are showed in lower case.

**Supplementary Table 5** Plasmids used in this study.

| Plasmids | Description | Reference |
|---|---|---|
| pYH7 | *ori(ColE1), ori(pIJ101), oriT(RK2), bla, tsr, aac(3)IV*, *E. coil-Streptomyces* shuttle vector | Sun et al. 2006 |
| pWHU2650 | pYH7 derivative with *Streptomyces* codon optimized *cas9* | Zeng et al. 2015 |
| pWHU2656 | pWHU2650+*actI-ORF2*-flanking sequences | Zeng et al. 2015 |
| pWHU2658 | pWUH2656+sgRNA targeting *actI-ORF2* | Zeng et al. 2015 |
| pWHU2739 | *E. coli* expression vector of Cas9, purificated Cas9 used to linearize pSCBE3 and pSCBE3-single to introduce HF mutations | Liu et al. 2015 |
| pUC57-SCBE-wocas9 | pUC57+*Streptomyces* codon optimized *cbe* without *cas9* | This work |
| pUC57-SABE-wocas9 | pUC57+*Streptomyces* codon optimized *abe* without *cas9* | This work |
| pUC57-MSCC | pUC57+multiple sgRNA cloning casette (MSCC) | This work |
| pUC57-SCBE2 | pUC57-SCBE-wocas9 +*Streptomyces* codon optimized *dcas9* | This work |
| pUC57-SCBE3 | pUC57-SCBE-wocas9 +*Streptomyces* codon optimized *ncas9* | This work |
| pUC57-SABEd | pUC57-SABE-wocas9 +*Streptomyces* codon optimized *dcas9* | This work |
| pUC57-SABEn | pUC57-SABE-wocas9 +*Streptomyces* codon optimized *ncas9* | This work |
| pYH7-wocos | pYH7 derivatve with *cos* site to be removed | This work |
| pYH7-MSCC | pYH7-wocos+multiple sgRNA cloning casette (MSCC) | This work |
| pSCBE2 | pYH7-MSCC+*Streptomyces* codon optimized *be2* (*tsr* replaced by *be2*) | This work |
| pSCBE3 | pYH7-MSCC+*Streptomyces* codon optimized *be3* (*tsr* replaced by *be3*) | This work |
| pSABEd | pYH7-MSCC+*Streptomyces* codon optimized *abed* (*tsr* replaced by *abed*) | This work |
| pSABEn | pYH7-MSCC+*Streptomyces* codon optimized *aben* (*tsr* replaced by *aben*) | This work |
| pSCBE2-single | pSCBE2-MSCC derivate for MSCC replaced by SSCC | This work |
| pSCBE3-single | pSCBE3-MSCC derivate for MSCC replaced by SSCC | This work |
| pSABEd-single | pSABEd-MSCC derivate for MSCC replaced by SSCC | This work |
| pSABEn-single | pSABEn-MSCC derivate for MSCC replaced by SSCC | This work |
| pSCBE2-*redD* | pSCBE2 derivate with *redD* spacer | This work |
| pSCBE3-*redD* | pSCBE3 derivate with *redD* spacer | This work |
| pSCBE2-*actI2* | pSCBE2 derivate with *actI-ORF2* spacer | This work |

| | | |
|---|---|---|
| pSCBE3-*actI2* | pSCBE3 derivate with *actI-ORF2* spacer | This work |
| pSCBE2-*redD-actI2* | pSCBE2 derivate with *redD* and *actI-ORF2* spacers | This work |
| pSCBE3-*redD-actI2* | pSCBE3 derivate with *redD* and *actI-ORF2* spacers | This work |
| pSCBE3-*pks11-1* | pSCBE3 derivate with *pks11-1* spacer | This work |
| pSCBE3-*pks11-2* | pSCBE3 derivate with *pks11-2* spacer | This work |
| pSCBE3-*pks11-3* | pSCBE3 derivate with *pks11-3* spacer | This work |
| pSCBE3-*pte-1* | pSCBE3 derivate with *pte-1* spacer | This work |
| pSCBE3-*pte-2* | pSCBE3 derivate with *pte-2* spacer | This work |
| pSCBE3-*pte-3* | pSCBE3 derivate with *pte-3* spacer | This work |
| pSCBE3-*pks2-1* | pSCBE3 derivate with *pks2-1* spacer | This work |
| pSCBE3-*pks2-2* | pSCBE3 derivate with *pks2-2* spacer | This work |
| pSCBE3-*pks2-3* | pSCBE3 derivate with *pks2-3* spacer | This work |
| pSCBE3-*pks3-1* | pSCBE3 derivate with *pks3-1* spacer | This work |
| pSCBE3-*pks3-2* | pSCBE3 derivate with *pks3-2* spacer | This work |
| pSCBE3-*pks3-3* | pSCBE3 derivate with *pks3-3* spacer | This work |
| pSCBE3-*pks5-1* | pSCBE3 derivate with *pks5-1* spacer | This work |
| pSCBE3-*pks5-2* | pSCBE3 derivate with *pks5-2* spacer | This work |
| pSCBE3-*pks5-3* | pSCBE3 derivate with *pks5-3* spacer | This work |
| pSCBE3-*pks9-1* | pSCBE3 derivate with *pks9-1* spacer | This work |
| pSCBE3-*pks9-2* | pSCBE3 derivate with *pks9-2* spacer | This work |
| pSCBE3-*pks9-3* | pSCBE3 derivate with *pks9-3* spacer | This work |
| pSCBE3-*olm-1* | pSCBE3 derivate with *olm-1* spacer | This work |
| pSCBE3-*olm-2* | pSCBE3 derivate with *olm-2* spacer | This work |
| pSCBE3-*olm-3* | pSCBE3 derivate with *olm-3* spacer | This work |
| pSCBE3-*pks8-1* | pSCBE3 derivate with *pks8-1* spacer | This work |
| pSCBE3-*pks8-2* | pSCBE3 derivate with *pks8-2* spacer | This work |
| pSCBE3-*pks8-3* | pSCBE3 derivate with *pks8-3* spacer | This work |
| pSCBE3-*rpp-1* | pSCBE3 derivate with *rpp-1* spacer | This work |
| pSCBE3-*rpp-2* | pSCBE3 derivate with *rpp-2* spacer | This work |
| pSCBE3-*rpp-3* | pSCBE3 derivate with *rpp-3* spacer | This work |
| pSCBE3-*pks4-1* | pSCBE3 derivate with *pks4-1* spacer | This work |
| pSCBE3-*pks4-2* | pSCBE3 derivate with *pks4-2* spacer | This work |

| | | |
|---|---|---|
| pSCBE3-*pks4-3* | pSCBE3 derivate with *pks4-3* spacer | This work |
| pSCBE3-*pks1-1* | pSCBE3 derivate with *pks1-1* spacer | This work |
| pSCBE3-*pks1-2* | pSCBE3 derivate with *pks1-2* spacer | This work |
| pSCBE3-*pks1-3* | pSCBE3 derivate with *pks1-3* spacer | This work |
| pSCBE3-5_sgRNA_array | pSCBE3 derivate with *olm-1, pks8-1, rpp-2, pks4-2* and *pks1-2* spacers | This work |
| pSCBE3-11_sgRNA_array | pSCBE3 derivate with *pks11-1, pte-3, pks2-2, pks3-1, pks5-1, pks9-2, olm-1, pks8-1, rpp-2, pks4-2* and *pks1-2* spacers | This work |
| pSCBE3-HF | pSCBE3 derivate with nCas9 replaced by HF-nCas9 | This work |
| pSCBE3-HF-single | pSCBE3-single derivative with nCas9 replaced by HF-nCas9 | This work |
| pSCBE3-HF-*olm* | pSCBE3-HF derivative with *olm* spacer | This work |
| pSABEd-*actVB* | pSABEd derivate with *actVB* spacer | This work |
| pSABEn-*actVB* | pSABEn derivate with *actVB* spacer | This work |

*ori(ColE1)*, *E. coli* replicon; *ori(pIJ101)*, *Streptomyces* replicon; *oriT(RK2)*, origin of transfer derived from RK2 plasmid; *bla*, ampicillin resistance gene; *tsr*, thiostrepton resistance gene; *aac(3)IV*, apramycin resistance gene; *cos*, cohesive end site.

**Supplementary Note 1** Complete sequence of the *Streptomyces* codon optimized base editors. Blue, APOBEC1; magenta, linkers; black, Cas9 variants; brown, UGI; green, TadA and TadA*; yellow highlight, Cas9 mutation in HNH domain (D10A); green highlight, Cas9 mutation in RuvC domain (H840A); cyan highlight, HF mutations (N497A, R661A, Q695A and Q926A).

> *be2*

ATGTCGTCCGAGACCGGCCCGGTGGCCGTGGACCCGACCCTGCGTCGCCGCATCGAGCCGCACGAGTTCGAGGTGTTCTTCGACCCGCGGGAG
CTGCGCAAGGAGACCTGCCTGCTGTACGAGATCAACTGGGGCGGCCGGCACTCGATCTGGCGCCACACCTCGCAGAACACCAACAAGCACGTC
GAGGTGAACTTCATCGAGAAGTTCACCACCGAGCGGTACTTCTGCCCGAACACCCGCTGCTCGATCACCTGGTTCCTGTCCTGGTCCCCGTGC
GGCGAGTGCTCCCGGGCGATCACCGAGTTCCTGTCCCGCTACCCGCACGTCACCCTGTTCATCTACATCGCCCGGCTGTACCACCACGCCGAC
CCGCGGAACCGGCAGGGCCTGCGCGACCTGATCTCCTCCGGCGTGACCATCCAGATCATGACCGAGCAGGAGTCCGGCTACTGCTGGCGGAAC
TTCGTCAACTACTCGCCGTCCAACGAGGCCCACTGGCCGCGGTACCCGCACCTGTGGGTGCGCCTGTACGTCCTGGAGCTGTACTGCATCATC
CTGGGCCTGCCGCCGTGCCTGAACATCCTGCGTCGCAAGCAGCCGCAGCTGACCTTCTTCACCATCGCCCTGCAGTCCTGCCACTACCAGCGC
CTGCCGCCGCACATCCTGTGGGCGACCGGCCTGAAGTCCGGCTCGGAGACCCCGGGCACCTCGGAGTCCGCCACCCCGGAGTCCGACAAGAAG
TACAGCATCGGCCTGGCCATCGGCACCAACAGCGTGGGCTGGGCGGTCATCACCGACGAGTACAAGGTCCCCTCCAAGAAGTTCAAGGTCCTG
GGCAACACCGACCGGCACTCGATCAAGAAGAACCTGATCGGCGCCCTGCTCTTCGACAGCGGCGAGACCGCCGAGGCGACCCGCCTGAAGCGG
ACCGCGCGCCGGCGCTACACCCGGCGCAAGAACCGCATCTGCTACCTCCAGGAAATCTTCTCCAACGAGATGGCCAAGGTCGACGACTCGTTC
TTCCACCGGCTCGAGGAGAGCTTCCTGGTGGAGGAGGACAAGAAGCACGAGCGCCACCCGATCTTCGGCAACATCGTCGACGAGGTGGCCTAC
CACGAGAAGTACCCCACCATCTACCACCTCCGCAAGAAGCTGGTCGACTCGACCGACAAGGCGGACCTGCGGCTCATCTACCTGGCCCTCGCG
CACATGATCAAGTTCCGCGGCCACTTCCTCATCGAGGGCGACCTGAACCCGGACAACTCCGACGTCGACAAGCTCTTCATCCAGCTGGTGCAG
ACCTACAACCAGCTGTTCGAGGAGAACCCCATCAACGCCAGCGGCGTCGACGCCAAGGCGATCCTCTCCGCGCGCCTGAGCAAGTCCCGGCGC
CTGGAGAACCTCATCGCCCAGCTGCCGGGCGAGAAGAAGAACGGCCTCTTCGGCAACCTGATCGCGCTGTCGCTCGGCCTGACCCCCAACTTC
AAGAGCAACTTCGACCTGGCCGAGGACGCGAAGCTCCAGCTGTCCAAGGACACCTACGACGACGACCTGGACAACCTGCTCGCCCAGATCGGC
GACCAGTACGCGGACCTCTTCCTGGCCGCGAAGAACCTCTCGGACGCCATCCTGCTCAGCGACATCCTGCGGGTCAACACCGAGATCACCAAG
GCCCCGCTGTCGGCGAGCATGATCAAGCGGTACGACGAGCACCACCAGGACCTGACCCTGCTCAAGGCCCTCGTGCGCCAGCAGCTGCCCGAG
AAGTACAAGGAAATCTTCTTCGACCAGTCCAAGAACGGCTACGCCGGCTACATCGACGGCGGCGCGTCGCAGGAGGAGTTCTACAAGTTCATC
AAGCCGATCCTGGAGAAGATGGACGGCACCGAGGAGCTGCTCGTCAAGCTGAACCGCGAGGACCTGCTCCGCAAGCAGCGGACCTTCGACAAC
GGCTCCATCCCGCACCAGATCCACCTGGGCGAGCTGCACGCCATCCTCCGGCGCCAGGAGGACTTCTACCCCTTCCTGAAGGACAACCGCGAG
AAGATCGAGAAGATCCTGACCTTCCGCATCCCGTACTACGTCGGCCCCCTGGCCCGCGGCAACTCCCGGTTCGCGTGGATGACCCGGAAGTCG
GAGGAGACCATCACCCCGTGGAACTTCGAGGAGGTCGTGGACAAGGGCGCCTCCGCCCAGTCGTTCATCGAGCGCATGACCAACTTCGACAAG
AACCTCCCGAACGAGAAGGTCCTGCCCAAGCACTCCCTGCTCTACGAGTACTTCACCGTGTACAACGAGCTGACCAAGGTCAAGTACGTGACC
GAGGGCATGCGGAAGCCGGCCTTCCTGTCGGGCGAGCAGAAGAAGGCGATCGTCGACCTGCTCTTCAAGACCAACCGCAAGGTCACCGTGAAG
CAGCTGAAGGAGGACTACTTCAAGAAGATCGAGTGCTTCGACTCCGTCGAGATCAGCGGCGTGGAGGACCGCTTCAACGCCTCCCTGGGCACC
TACCACGACCTGCTCAAGATCATCAAGGACAAGGACTTCCTCGACAACGAGGAGAACGAGGACATCCTGGAGGACATCGTCCTCACCCTGACC
CTCTTCGAGGACCGCGAGATGATCGAGGAGCGGCTCAAGACCTACGCCCACCTGTTCGACGACAAGGTGATGAAGCAGCTGAAGCGGCGCCGG
TACACCGGCTGGGGCCGCCTCTCCCGGAAGCTGATCAACGGCATCCGGGACAAGCAGAGCGGCAAGACCATCCTGGACTTCCTCAAGTCCGAC
GGCTTCGCCAACCGCAACTTCATGCAGCTCATCCACGACGACAGCCTGACCTTCAAGGAGGACATCCAGAAGGCCCAGGTCTCGGGCCAGGGC
GACAGCCTCCACGAGCACATCGCCAACCTGGCGGGCTCCCCGGCGATCAAGAAGGGCATCCTCCAGACCGTCAAGGTCGTGGACGAGCTGGTC
AAGGTGATGGGCCGCCACAAGCCCGAGAACATCGTGATCGAGATGGCCCGGGAGAACCAGACCACCCAGAAGGGCCAGAAGAACTCGCGCGAG
CGGATGAAGCGGATCGAGGAGGGCATCAAGGAGCTGGGCAGCCAGATCCTGAAGGAGCACCCGGTCGAGAACACCCAGCTCCAGAACGAGAAG
CTGTACCTCTACTACCTCCAGAACGGCCGCGACATGTACGTGGACCAGGAGCTGGACATCAACCGGCTGTCCGACTACGACGTCGACGCCATC
GTGCCGCAGTCCTTCCTGAAGGACGACTCGATCGACAACAAGGTCCTGACCCGCTCGGACAAGAACCGGGGCAAGTCCGACAACGTGCCCTCG
GAGGAGGTCGTGAAGAAGATGAAGAACTACTGGCGCCAGCTGCTCAACGCCAAGCTCATCACCCAGCGCAAGTTCGACAACCTGACCAAGGCC
GAGCGGGGCGGCCTGAGCGAGCTGGACAAGGCGGGCTTCATCAAGCGCCAGCTGGTCGAGACCCGGCAGATCACCAAGCACGTGGCCCAGATC
CTGGACTCCCGGATGAACACCAAGTACGACGAGAACGACAAGCTGATCCGCGAGGTCAAGGTGATCACCCTCAAGAGCAAGCTGGTCTCCGAC
TTCCGCAAGGACTTCCAGTTCTACAAGGTCCGGGAGATCAACAACTACCACCACGCCCACGACGCGTACCTGAACGCCGTCGTGGGCACCGCG
CTGATCAAGAAGTACCCGAAGCTGGAGTCCGAGTTCGTCTACGGCGACTACAAGGTCTACGACGTGCGCAAGATGATCGCCAAGAGCGAGCAG
GAGATCGGCAAGGCCACCGCGAAGTACTTCTTCTACTCCAACATCATGAACTTCTTCAAGACCGAGATCACCCTGGCCAACGGCGAGATCCGC
AAGCGGCCCCTGATCGAGACCAACGGCGAGACCGGCGAGATCGTCTGGGACAAGGGCCGCGACTTCGCCACCGTCCGGAAGGTGCTGTCGATG
CCGCAGGTCAACATCGTGAAGAAGACCGAGGTGCAGACCGGCGGCTTCAGCAAGGAGTCCATCCTCCCCAAGCGCAACAGCGACAAGCTGATC
GCCCGGAAGAAGGACTGGGACCCGAAGAAGTACGGCGGCTTCGACAGCCCCACCGTCGCCTACTCCGTGCTGGTCGTGGCGAAGGTCGAGAAG
GGCAAGAGCAAGAAGCTGAAGTCCGTGAAGGAGCTGCTCGGCATCACCATCATGGAGCGCTCCTCGTTCGAGAAGAACCCGATCGACTTCCTG
GAGGCCAAGGGCTACAAGGAGGTCAAGAAGGACCTCATCATCAAGCTGCCCAAGTACAGCCTGTTCGAGCTGGAGAACGGCCGCAAGCGGATG
CTCGCCTCCGCGGGCGAGCTGCAAAAGGGCAACGAGCTGGCCCTCCCGTCGAAGTACGTCAACTTCCTGTACCTCGCGTCCCACTACGAGAAG
CTGAAGGGCTCGCCCGAGGACAACGAGCAGAAGCAGCTCTTCGTGGAGCAGCACAAGCACTACCTGGACGAGATCATCGAGCAGATCAGCGAG
TTCAGCAAGCGCGTCATCCTGGCCGACGCGAACCTCGACAAGGTGCTGTCCGCCTACAACAAGCACCGCGACAAGCCGATCCGGGAGCAGGCG
GAGAACATCATCCACCTGTTCACCCTCACCAACCTGGGTGCCCCGGCCGCCTTCAAGTACTTCGACACCACCATCGACCGCAAGCGGTACACC
TCCACCAAGGAGGTCCTCGACGCGACCCTGATCCACCAGAGCATCACCGGCCTGTACGAGACCCGCATCGACCTGAGCCAGCTGGGCGGCGAC
TCGGGCGGCTCCACCAACCTGTCGGACATCATCGAGAAGGAGACCGGCAAGCAGCTGGTGATCCAGGAGTCCATCCTGATGCTGCCGGAGGAA
GTGGAGGAAGTGATCGGCAACAAGCCGGAGTCGGACATCCTGGTGCACACCGCCTACGACGAGTCGACCGACGAGAACGTCATGCTGCTGACC
TCGGACGCCCCGGAGTACAAGCCGTGGGCCCTGGTCATCCAGGACTCCAACGGCGAGAACAAGATCAAGATGCTGTGA

*>be3*

ATGTCGTCCGAGACCGGCCCGGTGGCCGTGGACCCGACCCTGCGTCGCCGCATCGAGCCGCACGAGTTCGAGGTGTTCTTCGACCCGCGGGAG
CTGCGCAAGGAGACCTGCCTGCTGTACGAGATCAACTGGGGCGGCCGGCACTCGATCTGGCGCCACACCTCGCAGAACACCAACAAGCACGTC
GAGGTGAACTTCATCGAGAAGTTCACCACCGAGCGGTACTTCTGCCCGAACACCCGCTGCTCGATCACCTGGTTCCTGTCCTGGTCCCCGTGC
GGCGAGTGCTCCCGGGCGATCACCGAGTTCCTGTCCCGCTACCCGCACGTCACCCTGTTCATCTACATCGCCCGGCTGTACCACCACGCCGAC
CCGCGGAACCGGCAGGGCCTGCGCGACCTGATCTCCTCCGGCGTGACCATCCAGATCATGACCGAGCAGGAGTCCGGCTACTGCTGGCGGAAC
TTCGTCAACTACTCGCCGTCCAACGAGGCCCACTGGCCGCGGTACCCGCACCTGTGGGTGCGCCTGTACGTCCTGGAGCTGTACTGCATCATC
CTGGGCCTGCCGCCGTGCCTGAACATCCTGCGTCGCAAGCAGCCGCAGCTGACCTTCTTCACCATCGCCCTGCAGTCCTGCCACTACCAGCGC
CTGCCGCCGCACATCCTGTGGGCGACCGGCCTGAAGTCCGGCTCGGAGACCCCGGGCACCTCGGAGTCCGCCACCCCGGAGTCCGACAAGAAG
TACAGCATCGGCCTGGCCATCGGCACCAACAGCGTGGGCTGGGCGGTCATCACCGACGAGTACAAGGTCCCCTCCAAGAAGTTCAAGGTCCTG
GGCAACACCGACCGGCACTCGATCAAGAAGAACCTGATCGGCGCCCTGCTCTTCGACAGCGGCGAGACCGCCGAGGCGACCCGCCTGAAGCGG
ACCGCGCGCCGGCGCTACACCCGGCGCAAGAACCGCATCTGCTACCTCCAGGAAATCTTCTCCAACGAGATGGCCAAGGTCGACGACTCGTTC
TTCCACCGGCTCGAGGAGAGCTTCCTGGTGGAGGAGGACAAGAAGCACGAGCGCCACCCGATCTTCGGCAACATCGTCGACGAGGTGGCCTAC
CACGAGAAGTACCCCACCATCTACCACCTCCGCAAGAAGCTGGTCGACTCGACCGACAAGGCGGACCTGCGGCTCATCTACCTGGCCCTCGCG
CACATGATCAAGTTCCGCGGCCACTTCCTCATCGAGGGCGACCTGAACCCGGACAACTCCGACGTCGACAAGCTCTTCATCCAGCTGGTGCAG
ACCTACAACCAGCTGTTCGAGGAGAACCCCATCAACGCCAGCGGCGTCGACGCCAAGGCGATCCTCTCCGCGCGCCTGAGCAAGTCCCGGCGC
CTGGAGAACCTCATCGCCCAGCTGCCGGGCGAGAAGAAGAACGGCCTCTTCGGCAACCTGATCGCGCTGTCGCTCGGCCTGACCCCCAACTTC
AAGAGCAACTTCGACCTGGCCGAGGACGCGAAGCTCCAGCTGTCCAAGGACACCTACGACGACGACCTGGACAACCTGCTCGCCCAGATCGGC
GACCAGTACGCGGACCTCTTCCTGGCCGCGAAGAACCTCTCGGACGCCATCCTGCTCAGCGACATCCTGCGGGTCAACACCGAGATCACCAAG
GCCCCGCTGTCGGCGAGCATGATCAAGCGGTACGACGAGCACCACCAGGACCTGACCCTGCTCAAGGCCCTCGTGCGCCAGCAGCTGCCCGAG
AAGTACAAGGAAATCTTCTTCGACCAGTCCAAGAACGGCTACGCCGGCTACATCGACGGCGGCGCGTCGCAGGAGGAGTTCTACAAGTTCATC
AAGCCGATCCTGGAGAAGATGGACGGCACCGAGGAGCTGCTCGTCAAGCTGAACCGCGAGGACCTGCTCCGCAAGCAGCGGACCTTCGACAAC
GGCTCCATCCCGCACCAGATCCACCTGGGCGAGCTGCACGCCATCCTCCGGCGCCAGGAGGACTTCTACCCCTTCCTGAAGGACAACCGCGAG
AAGATCGAGAAGATCCTGACCTTCCGCATCCCGTACTACGTCGGCCCCCTGGCCCGCGGCAACTCCCGGTTCGCGTGGATGACCCGGAAGTCG
GAGGAGACCATCACCCCGTGGAACTTCGAGGAGGTCGTGGACAAGGGCGCCTCCGCCCAGTCGTTCATCGAGCGCATGACCAACTTCGACAAG
AACCTCCCGAACGAGAAGGTCCTGCCCAAGCACTCCCTGCTCTACGAGTACTTCACCGTGTACAACGAGCTGACCAAGGTCAAGTACGTGACC
GAGGGCATGCGGAAGCCGGCCTTCCTGTCGGGCGAGCAGAAGAAGGCGATCGTCGACCTGCTCTTCAAGACCAACCGCAAGGTCACCGTGAAG
CAGCTGAAGGAGGACTACTTCAAGAAGATCGAGTGCTTCGACTCCGTCGAGATCAGCGGCGTGGAGGACCGCTTCAACGCCTCCCTGGGCACC
TACCACGACCTGCTCAAGATCATCAAGGACAAGGACTTCCTCGACAACGAGGAGAACGAGGACATCCTGGAGGACATCGTCCTCACCCTGACC
CTCTTCGAGGACCGCGAGATGATCGAGGAGCGGCTCAAGACCTACGCCCACCTGTTCGACGACAAGGTGATGAAGCAGCTGAAGCGGCGCCGG
TACACCGGCTGGGGCCGCCTCTCCCGGAAGCTGATCAACGGCATCCGGGACAAGCAGAGCGGCAAGACCATCCTGGACTTCCTCAAGTCCGAC
GGCTTCGCCAACCGCAACTTCATGCAGCTCATCCACGACGACAGCCTGACCTTCAAGGAGGACATCCAGAAGGCCCAGGTCTCGGGCCAGGGC
GACAGCCTCCACGAGCACATCGCCAACCTGGCGGGCTCCCCGGCGATCAAGAAGGGCATCCTCCAGACCGTCAAGGTCGTGGACGAGCTGGTC
AAGGTGATGGGCCGCCACAAGCCCGAGAACATCGTGATCGAGATGGCCCGGGAGAACCAGACCACCCAGAAGGGCCAGAAGAACTCGCGCGAG
CGGATGAAGCGGATCGAGGAGGGCATCAAGGAGCTGGGCAGCCAGATCCTGAAGGAGCACCCGGTCGAGAACACCCAGCTCCAGAACGAGAAG
CTGTACCTCTACTACCTCCAGAACGGCCGCGACATGTACGTGGACCAGGAGCTGGACATCAACCGGCTGTCCGACTACGACGTCGACCACATC
GTGCCGCAGTCCTTCCTGAAGGACGACTCGATCGACAACAAGGTCCTGACCCGCTCGGACAAGAACCGGGGCAAGTCCGACAACGTGCCCTCG
GAGGAGGTCGTGAAGAAGATGAAGAACTACTGGCGCCAGCTGCTCAACGCCAAGCTCATCACCCAGCGCAAGTTCGACAACCTGACCAAGGCC
GAGCGGGGCGGCCTGAGCGAGCTGGACAAGGCGGGCTTCATCAAGCGCCAGCTGGTCGAGACCCGGCAGATCACCAAGCACGTGGCCCAGATC
CTGGACTCCCGGATGAACACCAAGTACGACGAGAACGACAAGCTGATCCGCGAGGTCAAGGTGATCACCCTCAAGAGCAAGCTGGTCTCCGAC
TTCCGCAAGGACTTCCAGTTCTACAAGGTCCGGGAGATCAACAACTACCACCACGCCCACGACGCGTACCTGAACGCCGTCGTGGGCACCGCG
CTGATCAAGAAGTACCCGAAGCTGGAGTCCGAGTTCGTCTACGGCGACTACAAGGTCTACGACGTGCGCAAGATGATCGCCAAGAGCGAGCAG
GAGATCGGCAAGGCCACCGCGAAGTACTTCTTCTACTCCAACATCATGAACTTCTTCAAGACCGAGATCACCCTGGCCAACGGCGAGATCCGC
AAGCGGCCCCTGATCGAGACCAACGGCGAGACCGGCGAGATCGTCTGGGACAAGGGCCGCGACTTCGCCACCGTCCGGAAGGTGCTGTCGATG
CCGCAGGTCAACATCGTGAAGAAGACCGAGGTGCAGACCGGCGGCTTCAGCAAGGAGTCCATCCTCCCCAAGCGCAACAGCGACAAGCTGATC
GCCCGGAAGAAGGACTGGGACCCGAAGAAGTACGGCGGCTTCGACAGCCCCACCGTCGCCTACTCCGTGCTGGTCGTGGCGAAGGTCGAGAAG
GGCAAGAGCAAGAAGCTGAAGTCCGTGAAGGAGCTGCTCGGCATCACCATCATGGAGCGCTCCTCGTTCGAGAAGAACCCGATCGACTTCCTG
GAGGCCAAGGGCTACAAGGAGGTCAAGAAGGACCTCATCATCAAGCTGCCCAAGTACAGCCTGTTCGAGCTGGAGAACGGCCGCAAGCGGATG
CTCGCCTCCGCGGGCGAGCTGCAAAAGGGCAACGAGCTGGCCCTCCCGTCGAAGTACGTCAACTTCCTGTACCTCGCGTCCCACTACGAGAAG
CTGAAGGGCTCGCCCGAGGACAACGAGCAGAAGCAGCTCTTCGTGGAGCAGCACAAGCACTACCTGGACGAGATCATCGAGCAGATCAGCGAG
TTCAGCAAGCGCGTCATCCTGGCCGACGCGAACCTCGACAAGGTGCTGTCCGCCTACAACAAGCACCGCGACAAGCCGATCCGGGAGCAGGCG
GAGAACATCATCCACCTGTTCACCCTCACCAACCTGGGTGCCCCGGCCGCCTTCAAGTACTTCGACACCACCATCGACCGCAAGCGGTACACC
TCCACCAAGGAGGTCCTCGACGCGACCCTGATCCACCAGAGCATCACCGGCCTGTACGAGACCCGCATCGACCTGAGCCAGCTGGGCGGCGAC
TCGGGCGGCTCCACCAACCTGTCGGACATCATCGAGAAGGAGACCGGCAAGCAGCTGGTGATCCAGGAGTCCATCCTGATGCTGCCGGAGGAA
GTGGAGGAAGTGATCGGCAACAAGCCGGAGTCGGACATCCTGGTGCACACCGCCTACGACGAGTCGACCGACGAGAACGTCATGCTGCTGACC
TCGGACGCCCCGGAGTACAAGCCGTGGGCCCTGGTCATCCAGGACTCCAACGGCGAGAACAAGATCAAGATGCTGTGA

*>be3-HF*

ATGTCGTCCGAGACCGGCCCGGTGGCCGTGGACCCGACCCTGCGTCGCCGCATCGAGCCGCACGAGTTCGAGGTGTTCTTCGACCCGCGGGAG
CTGCGCAAGGAGACCTGCCTGCTGTACGAGATCAACTGGGGCGGCCGGCACTCGATCTGGCGCCACACCTCGCAGAACACCAACAAGCACGTC
GAGGTGAACTTCATCGAGAAGTTCACCACCGAGCGGTACTTCTGCCCGAACACCCGCTGCTCGATCACCTGGTTCCTGTCCTGGTCCCCGTGC
GGCGAGTGCTCCCGGGCGATCACCGAGTTCCTGTCCCGCTACCCGCACGTCACCCTGTTCATCTACATCGCCCGGCTGTACCACCACGCCGAC
CCGCGGAACCGGCAGGGCCTGCGCGACCTGATCTCCTCCGGCGTGACCATCCAGATCATGACCGAGCAGGAGTCCGGCTACTGCTGGCGGAAC
TTCGTCAACTACTCGCCGTCCAACGAGGCCCACTGGCCGCGGTACCCGCACCTGTGGGTGCGCCTGTACGTCCTGGAGCTGTACTGCATCATC
CTGGGCCTGCCGCCGTGCCTGAACATCCTGCGTCGCAAGCAGCCGCAGCTGACCTTCTTCACCATCGCCCTGCAGTCCTGCCACTACCAGCGC
CTGCCGCCGCACATCCTGTGGGCGACCGGCCTGAAGTCCGGCTCGGAGACCCCGGGCACCTCGGAGTCCGCCACCCCGGAGTCCGACAAGAAG
TACAGCATCGGCCTGGCCATCGGCACCAACAGCGTGGGCTGGGCGGTCATCACCGACGAGTACAAGGTCCCCTCCAAGAAGTTCAAGGTCCTG
GGCAACACCGACCGGCACTCGATCAAGAAGAACCTGATCGGCGCCCTGCTCTTCGACAGCGGCGAGACCGCCGAGGCGACCCGCCTGAAGCGG
ACCGCGCGCCGGCGCTACACCCGGCGCAAGAACCGCATCTGCTACCTCCAGGAAATCTTCTCCAACGAGATGGCCAAGGTCGACGACTCGTTC
TTCCACCGGCTCGAGGAGAGCTTCCTGGTGGAGGAGGACAAGAAGCACGAGCGCCACCCGATCTTCGGCAACATCGTCGACGAGGTGGCCTAC
CACGAGAAGTACCCCACCATCTACCACCTCCGCAAGAAGCTGGTCGACTCGACCGACAAGGCGGACCTGCGGCTCATCTACCTGGCCCTCGCG
CACATGATCAAGTTCCGCGGCCACTTCCTCATCGAGGGCGACCTGAACCCGGACAACTCCGACGTCGACAAGCTCTTCATCCAGCTGGTGCAG
ACCTACAACCAGCTGTTCGAGGAGAACCCCATCAACGCCAGCGGCGTCGACGCCAAGGCGATCCTCTCCGCGCGCCTGAGCAAGTCCCGGCGC
CTGGAGAACCTCATCGCCCAGCTGCCGGGCGAGAAGAAGAACGGCCTCTTCGGCAACCTGATCGCGCTGTCGCTCGGCCTGACCCCCAACTTC
AAGAGCAACTTCGACCTGGCCGAGGACGCGAAGCTCCAGCTGTCCAAGGACACCTACGACGACGACCTGGACAACCTGCTCGCCCAGATCGGC
GACCAGTACGCGGACCTCTTCCTGGCCGCGAAGAACCTCTCGGACGCCATCCTGCTCAGCGACATCCTGCGGGTCAACACCGAGATCACCAAG
GCCCCGCTGTCGGCGAGCATGATCAAGCGGTACGACGAGCACCACCAGGACCTGACCCTGCTCAAGGCCCTCGTGCGCCAGCAGCTGCCCGAG
AAGTACAAGGAAATCTTCTTCGACCAGTCCAAGAACGGCTACGCCGGCTACATCGACGGCGGCGCGTCGCAGGAGGAGTTCTACAAGTTCATC
AAGCCGATCCTGGAGAAGATGGACGGCACCGAGGAGCTGCTCGTCAAGCTGAACCGCGAGGACCTGCTCCGCAAGCAGCGGACCTTCGACAAC
GGCTCCATCCCGCACCAGATCCACCTGGGCGAGCTGCACGCCATCCTCCGGCGCCAGGAGGACTTCTACCCCTTCCTGAAGGACAACCGCGAG
AAGATCGAGAAGATCCTGACCTTCCGCATCCCGTACTACGTCGGCCCCCTGGCCCGCGGCAACTCCCGGTTCGCGTGGATGACCCGGAAGTCG
GAGGAGACCATCACCCCGTGGAACTTCGAGGAGGTCGTGGACAAGGGCGCCTCCGCCCAGTCGTTCATCGAGCGCATGACCGCCTTCGACAAG
AACCTCCCGAACGAGAAGGTCCTGCCCAAGCACTCCCTGCTCTACGAGTACTTCACCGTGTACAACGAGCTGACCAAGGTCAAGTACGTGACC
GAGGGCATGCGGAAGCCGGCCTTCCTGTCGGGCGAGCAGAAGAAGGCGATCGTCGACCTGCTCTTCAAGACCAACCGCAAGGTCACCGTGAAG
CAGCTGAAGGAGGACTACTTCAAGAAGATCGAGTGCTTCGACTCCGTCGAGATCAGCGGCGTGGAGGACCGCTTCAACGCCTCCCTGGGCACC
TACCACGACCTGCTCAAGATCATCAAGGACAAGGACTTCCTCGACAACGAGGAGAACGAGGACATCCTGGAGGACATCGTCCTCACCCTGACC
CTCTTCGAGGACCGCGAGATGATCGAGGAGCGGCTCAAGACCTACGCCCACCTGTTCGACGACAAGGTGATGAAGCAGCTGAAGCGGCGCCGG
TACACCGGCTGGGGCGCCCTCTCCCGGAAGCTGATCAACGGCATCCGGGACAAGCAGAGCGGCAAGACCATCCTGGACTTCCTCAAGTCCGAC
GGCTTCGCCAACCGCAACTTCATGGCGCTCATCCACGACGACAGCCTGACCTTCAAGGAGGACATCCAGAAGGCCCAGGTCTCGGGCCAGGGC
GACAGCCTCCACGAGCACATCGCCAACCTGGCGGGCTCCCCGGCGATCAAGAAGGGCATCCTCCAGACCGTCAAGGTCGTGGACGAGCTGGTC
AAGGTGATGGGCCGCCACAAGCCCGAGAACATCGTGATCGAGATGGCCCGGGAGAACCAGACCACCCAGAAGGGCCAGAAGAACTCGCGCGAG
CGGATGAAGCGGATCGAGGAGGGCATCAAGGAGCTGGGCAGCCAGATCCTGAAGGAGCACCCGGTCGAGAACACCCAGCTCCAGAACGAGAAG
CTGTACCTCTACTACCTCCAGAACGGCCGCGACATGTACGTGGACCAGGAGCTGGACATCAACCGGCTGTCCGACTACGACGTCGACCACATC
GTGCCGCAGTCCTTCCTGAAGGACGACTCGATCGACAACAAGGTCCTGACCCGCTCGGACAAGAACCGGGGCAAGTCCGACAACGTGCCCTCG
GAGGAGGTCGTGAAGAAGATGAAGAACTACTGGCGCCAGCTGCTCAACGCCAAGCTCATCACCCAGCGCAAGTTCGACAACCTGACCAAGGCC
GAGCGGGGCGGCCTGAGCGAGCTGGACAAGGCGGGCTTCATCAAGCGCCAGCTGGTCGAGACCCGGGCGATCACCAAGCACGTGGCCCAGATC
CTGGACTCCCGGATGAACACCAAGTACGACGAGAACGACAAGCTGATCCGCGAGGTCAAGGTGATCACCCTCAAGAGCAAGCTGGTCTCCGAC
TTCCGCAAGGACTTCCAGTTCTACAAGGTCCGGGAGATCAACAACTACCACCACGCCCACGACGCGTACCTGAACGCCGTCGTGGGCACCGCG
CTGATCAAGAAGTACCCGAAGCTGGAGTCCGAGTTCGTCTACGGCGACTACAAGGTCTACGACGTGCGCAAGATGATCGCCAAGAGCGAGCAG
GAGATCGGCAAGGCCACCGCGAAGTACTTCTTCTACTCCAACATCATGAACTTCTTCAAGACCGAGATCACCCTGGCCAACGGCGAGATCCGC
AAGCGGCCCCTGATCGAGACCAACGGCGAGACCGGCGAGATCGTCTGGGACAAGGGCCGCGACTTCGCCACCGTCCGGAAGGTGCTGTCGATG
CCGCAGGTCAACATCGTGAAGAAGACCGAGGTGCAGACCGGCGGCTTCAGCAAGGAGTCCATCCTCCCCAAGCGCAACAGCGACAAGCTGATC
GCCCGGAAGAAGGACTGGGACCCGAAGAAGTACGGCGGCTTCGACAGCCCCACCGTCGCCTACTCCGTGCTGGTCGTGGCGAAGGTCGAGAAG
GGCAAGAGCAAGAAGCTGAAGTCCGTGAAGGAGCTGCTCGGCATCACCATCATGGAGCGCTCCTCGTTCGAGAAGAACCCGATCGACTTCCTG
GAGGCCAAGGGCTACAAGGAGGTCAAGAAGGACCTCATCATCAAGCTGCCCAAGTACAGCCTGTTCGAGCTGGAGAACGGCCGCAAGCGGATG
CTCGCCTCCGCGGGCGAGCTGCAAAAGGGCAACGAGCTGGCCCTCCCGTCGAAGTACGTCAACTTCCTGTACCTCGCGTCCCACTACGAGAAG
CTGAAGGGCTCGCCCGAGGACAACGAGCAGAAGCAGCTCTTCGTGGAGCAGCACAAGCACTACCTGGACGAGATCATCGAGCAGATCAGCGAG
TTCAGCAAGCGCGTCATCCTGGCCGACGCGAACCTCGACAAGGTGCTGTCCGCCTACAACAAGCACCGCGACAAGCCGATCCGGGAGCAGGCG
GAGAACATCATCCACCTGTTCACCCTCACCAACCTGGGTGCCCCGGCCGCCTTCAAGTACTTCGACACCACCATCGACCGCAAGCGGTACACC
TCCACCAAGGAGGTCCTCGACGCGACCCTGATCCACCAGAGCATCACCGGCCTGTACGAGACCCGCATCGACCTGAGCCAGCTGGGCGGCGAC
TCGGGCGGCTCCACCAACCTGTCGGACATCATCGAGAAGGAGACCGGCAAGCAGCTGGTGATCCAGGAGTCCATCCTGATGCTGCCGGAGGAA
GTGGAGGAAGTGATCGGCAACAAGCCGGAGTCGGACATCCTGGTGCACACCGCCTACGACGAGTCGACCGACGAGAACGTCATGCTGCTGACC
TCGGACGCCCCGGAGTACAAGCCGTGGGCCCTGGTCATCCAGGACTCCAACGGCGAGAACAAGATCAAGATGCTGTGA

*>abed*

ATGTCCGAGGTGGAGTTCTCCCACGAGTACTGGATGCGCCACGCCCTGACCCTGGCCAAGCGCGCGTGGGACGAGCGCGAGGTGCCGGTGGGC
GCCGTGCTGGTCCACAACAACCGCGTGATCGGCGAGGGCTGGAACCGGCCGATCGGCCGGCACGACCCGACCGCCCACGCCGAGATCATGGCC
CTGCGCCAGGGCGGCCTGGTCATGCAGAACTACCGGCTGATCGACGCGACCCTGTACGTGACCCTGGAGCCGTGCGTCATGTGCGCGGGCGCC
ATGATCCACTCCCGCATCGGCCGGGTGGTGTTCGGCGCCCGGGACGCCAAGACCGGCGCCGCGGGCTCCCTGATGGACGTGCTGCACCACCCG
GGCATGAACCACCGCGTCGAGATCACCGAGGGCATCCTGGCGGACGAGTGCGCCGCGCTGCTGTCGGACTTCTTCCGGATGCGCCGGCAGGAG
ATCAAGGCCCAGAAGAAGGCGCAGTCGTCCACCGACTCGGGCGGCTCGTCCGGCGGCTCGTCCGGCTCCGAGACCCCGGGCACCTCGGAGTCC
GCGACCCCGGAGTCGTCCGGCGGCTCGTCCGGCGGCTCGTCCGAGGTCGAATTTTCCCATGAATACTGGATGCGGCACGCGCTGACCCTGGCC
AAGCGGGCCCGCGACGAGCGCGAGGTGCCCGTCGGCGCGGTGCTGGTCCTGAACAACCGGGTCATTGGTGAGGGCTGGAACCGCGCCATCGGC
CTGCACGACCCCACCGCGCATGCCGAAATCATGGCCCTCCGGCAGGGCGGCCTGGTGATGCAGAACTACCGCCTCATCGACGCCACCCTGTAC
GTGACCTTCGAGCCCTGCGTGATGTGTGCGGGTGCCATGATCCACTCCCGGATCGGCCGCGTCGTGTTCGGCGTGCGCAACGCCAAGACCGGC
GCGGCCGGCTCCCTGATGGACGTCCTGCACTACCCCGGCATGAACCACCGCGTGGAAATTACCGAGGGCATCCTGGCCGACGAGTGCGCGGCC
CTGCTGTGCTACTTCTTCCGCATGCCGCGGCAGGTGTTCAACGCGCAGAAGAAGGCGCAGTCGTCCACGGATTCGGGCGGCTCGTCCGGCGGC
TCGTCCGGGTCGGAAACCCCGGGTACCTCGGAGTCGGCCACCCCGGAGTCGTCCGGCGGCTCGTCCGGCGGCTCGGACAAGAAGTACAGCATC
GGCCTGGCCATCGGCACCAACAGCGTGGGCTGGGCGGTCATCACCGACGAGTACAAGGTCCCCTCCAAGAAGTTCAAGGTCCTGGGCAACACC
GACCGGCACTCGATCAAGAAGAACCTGATCGGCGCCCTGCTCTTCGACAGCGGCGAGACCGCCGAGGCGACCCGCCTGAAGCGGACCGCGCGC
CGGCGCTACACCCGGCGCAAGAACCGCATCTGCTACCTCCAGGAAATCTTCTCCAACGAGATGGCCAAGGTCGACGACTCGTTCTTCCACCGG
CTCGAGGAGAGCTTCCTGGTGGAGGAGGACAAGAAGCACGAGCGCCACCCGATCTTCGGCAACATCGTCGACGAGGTGGCCTACCACGAGAAG
TACCCCACCATCTACCACCTCCGCAAGAAGCTGGTCGACTCGACCGACAAGGCGGACCTGCGGCTCATCTACCTGGCCCTCGCGCACATGATC
AAGTTCCGCGGCCACTTCCTCATCGAGGGCGACCTGAACCCGGACAACTCCGACGTCGACAAGCTCTTCATCCAGCTGGTGCAGACCTACAAC
CAGCTGTTCGAGGAGAACCCCATCAACGCCAGCGGCGTCGACGCCAAGGCGATCCTCTCCGCGCGCCTGAGCAAGTCCCGGCGCCTGGAGAAC
CTCATCGCCCAGCTGCCGGGCGAGAAGAAGAACGGCCTCTTCGGCAACCTGATCGCGCTGTCGCTCGGCCTGACCCCCAACTTCAAGAGCAAC
TTCGACCTGGCCGAGGACGCGAAGCTCCAGCTGTCCAAGGACACCTACGACGACGACCTGGACAACCTGCTCGCCCAGATCGGCGACCAGTAC
GCGGACCTCTTCCTGGCCGCGAAGAACCTCTCGGACGCCATCCTGCTCAGCGACATCCTGCGGGTCAACACCGAGATCACCAAGGCCCCGCTG
TCGGCGAGCATGATCAAGCGGTACGACGAGCACCACCAGGACCTGACCCTGCTCAAGGCCCTCGTGCGCCAGCAGCTGCCCGAGAAGTACAAG
GAAATCTTCTTCGACCAGTCCAAGAACGGCTACGCCGGCTACATCGACGGCGGCGCGTCGCAGGAGGAGTTCTACAAGTTCATCAAGCCGATC
CTGGAGAAGATGGACGGCACCGAGGAGCTGCTCGTCAAGCTGAACCGCGAGGACCTGCTCCGCAAGCAGCGGACCTTCGACAACGGCTCCATC
CCGCACCAGATCCACCTGGGCGAGCTGCACGCCATCCTCCGGCGCCAGGAGGACTTCTACCCCTTCCTGAAGGACAACCGCGAGAAGATCGAG
AAGATCCTGACCTTCCGCATCCCGTACTACGTCGGCCCCCTGGCCCGCGGCAACTCCCGGTTCGCGTGGATGACCCGGAAGTCGGAGGAGACC
ATCACCCCGTGGAACTTCGAGGAGGTCGTGGACAAGGGCGCCTCCGCCCAGTCGTTCATCGAGCGCATGACCAACTTCGACAAGAACCTCCCG
AACGAGAAGGTCCTGCCCAAGCACTCCCTGCTCTACGAGTACTTCACCGTGTACAACGAGCTGACCAAGGTCAAGTACGTGACCGAGGGCATG
CGGAAGCCGGCCTTCCTGTCGGGCGAGCAGAAGAAGGCGATCGTCGACCTGCTCTTCAAGACCAACCGCAAGGTCACCGTGAAGCAGCTGAAG
GAGGACTACTTCAAGAAGATCGAGTGCTTCGACTCCGTCGAGATCAGCGGCGTGGAGGACCGCTTCAACGCCTCCCTGGGCACCTACCACGAC
CTGCTCAAGATCATCAAGGACAAGGACTTCCTCGACAACGAGGAGAACGAGGACATCCTGGAGGACATCGTCCTCACCCTGACCCTCTTCGAG
GACCGCGAGATGATCGAGGAGCGGCTCAAGACCTACGCCCACCTGTTCGACGACAAGGTGATGAAGCAGCTGAAGCGGCGCCGGTACACCGGC
TGGGGCCGCCTCTCCCGGAAGCTGATCAACGGCATCCGGGACAAGCAGAGCGGCAAGACCATCCTGGACTTCCTCAAGTCCGACGGCTTCGCC
AACCGCAACTTCATGCAGCTCATCCACGACGACAGCCTGACCTTCAAGGAGGACATCCAGAAGGCCCAGGTCTCGGGCCAGGGCGACAGCCTC
CACGAGCACATCGCCAACCTGGCGGGCTCCCCGGCGATCAAGAAGGGCATCCTCCAGACCGTCAAGGTCGTGGACGAGCTGGTCAAGGTGATG
GGCCGCCACAAGCCCGAGAACATCGTGATCGAGATGGCCCGGGAGAACCAGACCACCCAGAAGGGCCAGAAGAACTCGCGCGAGCGGATGAAG
CGGATCGAGGAGGGCATCAAGGAGCTGGGCAGCCAGATCCTGAAGGAGCACCCGGTCGAGAACACCCAGCTCCAGAACGAGAAGCTGTACCTC
TACTACCTCCAGAACGGCCGCGACATGTACGTGGACCAGGAGCTGGACATCAACCGGCTGTCCGACTACGACGTCGACGCCATCGTGCCGCAG
TCCTTCCTGAAGGACGACTCGATCGACAACAAGGTCCTGACCCGCTCGGACAAGAACCGGGGCAAGTCCGACAACGTGCCCTCGGAGGAGGTC
GTGAAGAAGATGAAGAACTACTGGCGCCAGCTGCTCAACGCCAAGCTCATCACCCAGCGCAAGTTCGACAACCTGACCAAGGCCGAGCGGGGC
GGCCTGAGCGAGCTGGACAAGGCGGGCTTCATCAAGCGCCAGCTGGTCGAGACCCGGCAGATCACCAAGCACGTGGCCCAGATCCTGGACTCC
CGGATGAACACCAAGTACGACGAGAACGACAAGCTGATCCGCGAGGTCAAGGTGATCACCCTCAAGAGCAAGCTGGTCTCCGACTTCCGCAAG
GACTTCCAGTTCTACAAGGTCCGGGAGATCAACAACTACCACCACGCCCACGACGCGTACCTGAACGCCGTCGTGGGCACCGCGCTGATCAAG
AAGTACCCGAAGCTGGAGTCCGAGTTCGTCTACGGCGACTACAAGGTCTACGACGTGCGCAAGATGATCGCCAAGAGCGAGCAGGAGATCGGC
AAGGCCACCGCGAAGTACTTCTTCTACTCCAACATCATGAACTTCTTCAAGACCGAGATCACCCTGGCCAACGGCGAGATCCGCAAGCGGCCC
CTGATCGAGACCAACGGCGAGACCGGCGAGATCGTCTGGGACAAGGGCCGCGACTTCGCCACCGTCCGGAAGGTGCTGTCGATGCCGCAGGTC
AACATCGTGAAGAAGACCGAGGTGCAGACCGGCGGCTTCAGCAAGGAGTCCATCCTCCCCAAGCGCAACAGCGACAAGCTGATCGCCCGGAAG
AAGGACTGGGACCCGAAGAAGTACGGCGGCTTCGACAGCCCCACCGTCGCCTACTCCGTGCTGGTCGTGGCGAAGGTCGAGAAGGGCAAGAGC
AAGAAGCTGAAGTCCGTGAAGGAGCTGCTCGGCATCACCATCATGGAGCGCTCCTCGTTCGAGAAGAACCCGATCGACTTCCTGGAGGCCAAG
GGCTACAAGGAGGTCAAGAAGGACCTCATCATCAAGCTGCCCAAGTACAGCCTGTTCGAGCTGGAGAACGGCCGCAAGCGGATGCTCGCCTCC
GCGGGCGAGCTGCAAAAGGGCAACGAGCTGGCCCTCCCGTCGAAGTACGTCAACTTCCTGTACCTCGCGTCCCACTACGAGAAGCTGAAGGGC
TCGCCCGAGGACAACGAGCAGAAGCAGCTCTTCGTGGAGCAGCACAAGCACTACCTGGACGAGATCATCGAGCAGATCAGCGAGTTCAGCAAG
CGCGTCATCCTGGCCGACGCGAACCTCGACAAGGTGCTGTCCGCCTACAACAAGCACCGCGACAAGCCGATCCGGGAGCAGGCGGAGAACATC
ATCCACCTGTTCACCCTCACCAACCTGGGTGCCCCGGCCGCCTTCAAGTACTTCGACACCACCATCGACCGCAAGCGGTACACCTCCACCAAG
GAGGTCCTCGACGCGACCCTGATCCACCAGAGCATCACCGGCCTGTACGAGACCCGCATCGACCTGAGCCAGCTGGGCGGCGACTGA

*>aben*

ATGTCCGAGGTGGAGTTCTCCCACGAGTACTGGATGCGCCACGCCCTGACCCTGGCCAAGCGCGCGTGGGACGAGCGCGAGGTGCCGGTGGGC
GCCGTGCTGGTCCACAACAACCGCGTGATCGGCGAGGGCTGGAACCGGCCGATCGGCCGGCACGACCCGACCGCCCACGCCGAGATCATGGCC
CTGCGCCAGGGCGGCCTGGTCATGCAGAACTACCGGCTGATCGACGCGACCCTGTACGTGACCCTGGAGCCGTGCGTCATGTGCGCGGGCGCC
ATGATCCACTCCCGCATCGGCCGGGTGGTGTTCGGCGCCCGGGACGCCAAGACCGGCGCCGCGGGCTCCCTGATGGACGTGCTGCACCACCCG
GGCATGAACCACCGCGTCGAGATCACCGAGGGCATCCTGGCGGACGAGTGCGCCGCGCTGCTGTCGGACTTCTTCCGGATGCGCCGGCAGGAG
ATCAAGGCCCAGAAGAAGGCGCAGTCGTCCACCGACTCGGGCGGCTCGTCCGGCGGCTCGTCCGGCTCCGAGACCCCGGGCACCTCGGAGTCC
GCGACCCCGGAGTCGTCCGGCGGCTCGTCCGGCGGCTCGTCCGAGGTCGAATTTTCCCATGAATACTGGATGCGGCACGCGCTGACCCTGGCC
AAGCGGGCCCGCGACGAGCGCGAGGTGCCCGTCGGCGCGGTGCTGGTCCTGAACAACCGGGTCATTGGTGAGGGCTGGAACCGCGCCATCGGC
CTGCACGACCCCACCGCGCATGCCGAAATCATGGCCCTCCGGCAGGGCGGCCTGGTGATGCAGAACTACCGCCTCATCGACGCCACCCTGTAC
GTGACCTTCGAGCCCTGCGTGATGTGTGCGGGTGCCATGATCCACTCCCGGATCGGCCGCGTCGTGTTCGGCGTGCGCAACGCCAAGACCGGC
GCGGCCGGCTCCCTGATGGACGTCCTGCACTACCCCGGCATGAACCACCGCGTGGAAATTACCGAGGGCATCCTGGCCGACGAGTGCGCGGCC
CTGCTGTGCTACTTCTTCCGCATGCCGCGGCAGGTGTTCAACGCGCAGAAGAAGGCGCAGTCGTCCACGGATTCGGGCGGCTCGTCCGGCGGC
TCGTCCGGGTCGGAAACCCCGGGTACCTCGGAGTCGGCCACCCCGGAGTCGTCCGGCGGCTCGTCCGGCGGCTCGGACAAGAAGTACAGCATC
GGCCTGGCCATCGGCACCAACAGCGTGGGCTGGGCGGTCATCACCGACGAGTACAAGGTCCCCTCCAAGAAGTTCAAGGTCCTGGGCAACACC
GACCGGCACTCGATCAAGAAGAACCTGATCGGCGCCCTGCTCTTCGACAGCGGCGAGACCGCCGAGGCGACCCGCCTGAAGCGGACCGCGCGC
CGGCGCTACACCCGGCGCAAGAACCGCATCTGCTACCTCCAGGAAATCTTCTCCAACGAGATGGCCAAGGTCGACGACTCGTTCTTCCACCGG
CTCGAGGAGAGCTTCCTGGTGGAGGAGGACAAGAAGCACGAGCGCCACCCGATCTTCGGCAACATCGTCGACGAGGTGGCCTACCACGAGAAG
TACCCCACCATCTACCACCTCCGCAAGAAGCTGGTCGACTCGACCGACAAGGCGGACCTGCGGCTCATCTACCTGGCCCTCGCGCACATGATC
AAGTTCCGCGGCCACTTCCTCATCGAGGGCGACCTGAACCCGGACAACTCCGACGTCGACAAGCTCTTCATCCAGCTGGTGCAGACCTACAAC
CAGCTGTTCGAGGAGAACCCCATCAACGCCAGCGGCGTCGACGCCAAGGCGATCCTCTCCGCGCGCCTGAGCAAGTCCCGGCGCCTGGAGAAC
CTCATCGCCCAGCTGCCGGGCGAGAAGAAGAACGGCCTCTTCGGCAACCTGATCGCGCTGTCGCTCGGCCTGACCCCCAACTTCAAGAGCAAC
TTCGACCTGGCCGAGGACGCGAAGCTCCAGCTGTCCAAGGACACCTACGACGACGACCTGGACAACCTGCTCGCCCAGATCGGCGACCAGTAC
GCGGACCTCTTCCTGGCCGCGAAGAACCTCTCGGACGCCATCCTGCTCAGCGACATCCTGCGGGTCAACACCGAGATCACCAAGGCCCCGCTG
TCGGCGAGCATGATCAAGCGGTACGACGAGCACCACCAGGACCTGACCCTGCTCAAGGCCCTCGTGCGCCAGCAGCTGCCCGAGAAGTACAAG
GAAATCTTCTTCGACCAGTCCAAGAACGGCTACGCCGGCTACATCGACGGCGGCGCGTCGCAGGAGGAGTTCTACAAGTTCATCAAGCCGATC
CTGGAGAAGATGGACGGCACCGAGGAGCTGCTCGTCAAGCTGAACCGCGAGGACCTGCTCCGCAAGCAGCGGACCTTCGACAACGGCTCCATC
CCGCACCAGATCCACCTGGGCGAGCTGCACGCCATCCTCCGGCGCCAGGAGGACTTCTACCCCTTCCTGAAGGACAACCGCGAGAAGATCGAG
AAGATCCTGACCTTCCGCATCCCGTACTACGTCGGCCCCCTGGCCCGCGGCAACTCCCGGTTCGCGTGGATGACCCGGAAGTCGGAGGAGACC
ATCACCCCGTGGAACTTCGAGGAGGTCGTGGACAAGGGCGCCTCCGCCCAGTCGTTCATCGAGCGCATGACCAACTTCGACAAGAACCTCCCG
AACGAGAAGGTCCTGCCCAAGCACTCCCTGCTCTACGAGTACTTCACCGTGTACAACGAGCTGACCAAGGTCAAGTACGTGACCGAGGGCATG
CGGAAGCCGGCCTTCCTGTCGGGCGAGCAGAAGAAGGCGATCGTCGACCTGCTCTTCAAGACCAACCGCAAGGTCACCGTGAAGCAGCTGAAG
GAGGACTACTTCAAGAAGATCGAGTGCTTCGACTCCGTCGAGATCAGCGGCGTGGAGGACCGCTTCAACGCCTCCCTGGGCACCTACCACGAC
CTGCTCAAGATCATCAAGGACAAGGACTTCCTCGACAACGAGGAGAACGAGGACATCCTGGAGGACATCGTCCTCACCCTGACCCTCTTCGAG
GACCGCGAGATGATCGAGGAGCGGCTCAAGACCTACGCCCACCTGTTCGACGACAAGGTGATGAAGCAGCTGAAGCGGCGCCGGTACACCGGC
TGGGGCCGCCTCTCCCGGAAGCTGATCAACGGCATCCGGGACAAGCAGAGCGGCAAGACCATCCTGGACTTCCTCAAGTCCGACGGCTTCGCC
AACCGCAACTTCATGCAGCTCATCCACGACGACAGCCTGACCTTCAAGGAGGACATCCAGAAGGCCCAGGTCTCGGGCCAGGGCGACAGCCTC
CACGAGCACATCGCCAACCTGGCGGGCTCCCCGGCGATCAAGAAGGGCATCCTCCAGACCGTCAAGGTCGTGGACGAGCTGGTCAAGGTGATG
GGCCGCCACAAGCCCGAGAACATCGTGATCGAGATGGCCCGGGAGAACCAGACCACCCAGAAGGGCCAGAAGAACTCGCGCGAGCGGATGAAG
CGGATCGAGGAGGGCATCAAGGAGCTGGGCAGCCAGATCCTGAAGGAGCACCCGGTCGAGAACACCCAGCTCCAGAACGAGAAGCTGTACCTC
TACTACCTCCAGAACGGCCGCGACATGTACGTGGACCAGGAGCTGGACATCAACCGGCTGTCCGACTACGACGTCGACCACATCGTGCCGCAG
TCCTTCCTGAAGGACGACTCGATCGACAACAAGGTCCTGACCCGCTCGGACAAGAACCGGGGCAAGTCCGACAACGTGCCCTCGGAGGAGGTC
GTGAAGAAGATGAAGAACTACTGGCGCCAGCTGCTCAACGCCAAGCTCATCACCCAGCGCAAGTTCGACAACCTGACCAAGGCCGAGCGGGGC
GGCCTGAGCGAGCTGGACAAGGCGGGCTTCATCAAGCGCCAGCTGGTCGAGACCCGGCAGATCACCAAGCACGTGGCCCAGATCCTGGACTCC
CGGATGAACACCAAGTACGACGAGAACGACAAGCTGATCCGCGAGGTCAAGGTGATCACCCTCAAGAGCAAGCTGGTCTCCGACTTCCGCAAG
GACTTCCAGTTCTACAAGGTCCGGGAGATCAACAACTACCACCACGCCCACGACGCGTACCTGAACGCCGTCGTGGGCACCGCGCTGATCAAG
AAGTACCCGAAGCTGGAGTCCGAGTTCGTCTACGGCGACTACAAGGTCTACGACGTGCGCAAGATGATCGCCAAGAGCGAGCAGGAGATCGGC
AAGGCCACCGCGAAGTACTTCTTCTACTCCAACATCATGAACTTCTTCAAGACCGAGATCACCCTGGCCAACGGCGAGATCCGCAAGCGGCCC
CTGATCGAGACCAACGGCGAGACCGGCGAGATCGTCTGGGACAAGGGCCGCGACTTCGCCACCGTCCGGAAGGTGCTGTCGATGCCGCAGGTC
AACATCGTGAAGAAGACCGAGGTGCAGACCGGCGGCTTCAGCAAGGAGTCCATCCTCCCCAAGCGCAACAGCGACAAGCTGATCGCCCGGAAG
AAGGACTGGGACCCGAAGAAGTACGGCGGCTTCGACAGCCCCACCGTCGCCTACTCCGTGCTGGTCGTGGCGAAGGTCGAGAAGGGCAAGAGC
AAGAAGCTGAAGTCCGTGAAGGAGCTGCTCGGCATCACCATCATGGAGCGCTCCTCGTTCGAGAAGAACCCGATCGACTTCCTGGAGGCCAAG
GGCTACAAGGAGGTCAAGAAGGACCTCATCATCAAGCTGCCCAAGTACAGCCTGTTCGAGCTGGAGAACGGCCGCAAGCGGATGCTCGCCTCC
GCGGGCGAGCTGCAAAAGGGCAACGAGCTGGCCCTCCCGTCGAAGTACGTCAACTTCCTGTACCTCGCGTCCCACTACGAGAAGCTGAAGGGC
TCGCCCGAGGACAACGAGCAGAAGCAGCTCTTCGTGGAGCAGCACAAGCACTACCTGGACGAGATCATCGAGCAGATCAGCGAGTTCAGCAAG
CGCGTCATCCTGGCCGACGCGAACCTCGACAAGGTGCTGTCCGCCTACAACAAGCACCGCGACAAGCCGATCCGGGAGCAGGCGGAGAACATC
ATCCACCTGTTCACCCTCACCAACCTGGGTGCCCCGGCCGCCTTCAAGTACTTCGACACCACCATCGACCGCAAGCGGTACACCTCCACCAAG
GAGGTCCTCGACGCGACCCTGATCCACCAGAGCATCACCGGCCTGTACGAGACCCGCATCGACCTGAGCCAGCTGGGCGGCGACTGA

**References**


1. Billon, P. *et al.* CRISPR-mediated base editing enables efficient disruption of eukaryotic genes through induction of STOP codons. *Mol. Cell* **67**, 1068-1079.e4 (2017).

2. Sun, Y. *et al.* Organization of the biosynthetic gene cluster in *Streptomyces* sp. DSM 4137 for the novel neuroprotectant polyketide meridamycin. *Microbiology* **152**, 3507–3515 (2006).

3. Zeng, H. *et al.* Highly efficient editing of the actinorhodin polyketide chain length factor gene in *Streptomyces coelicolor* M145 using CRISPR/Cas9-CodA(sm) combined system. *Appl. Microbiol. Biotechnol.* **99**, 10575–10585 (2015).

4. Liu, Y. *et al.* *In vitro* CRISPR/Cas9 system for efficient targeted DNA editing. *mBio* **6**, e01714–15 (2015).
